# The deSUMOylase SENP2 coordinates homologous recombination and non-homologous end joining by independent mechanisms

**DOI:** 10.1101/473991

**Authors:** Alexander J. Garvin, Alexandra K. Walker, Ruth M. Densham, Anoop Singh Chauhan, Helen R. Stone, Hannah L. Mackay, Mohammed Jamshad, Katarzyna Starowicz, Manuel Daza-Martin, Joanna R. Morris

**Affiliations:** Birmingham Centre for Genome Biology and Institute of Cancer and Genomic Sciences, University of Birmingham, Edgbaston, Birmingham, B15 2TT, United Kingdom

## Abstract

SUMOylation in the DNA double-strand break (DSB) response regulates recruitment, activity and clearance of repair factors. However, our understanding of a role for deSUMOylation in this process is limited. Here we identify different mechanistic roles for deSUMOylation in homologous recombination (HR) and non-homologous enjoining (NHEJ) through the investigation of the deSUMOylase SENP2. We find regulated deSUMOylation of MDC1 prevents excessive SUMOylation and its RNF4-VCP mediated clearance from DSBs, thereby promoting NHEJ. In contrast we show HR is differentially sensitive to SUMO availability and SENP2 activity is needed to provide SUMO. *SENP2* is amplified as part of the chromosome 3q amplification in many cancers. Increased SENP2 expression prolongs MDC1 foci retention and increases NHEJ and radioresistance. Collectively our data reveal that deSUMOylation differentially primes cells for responding to DSBs and demonstrates the ability of SENP2 to tune DSB repair responses.

## Introduction

The cellular response to DNA double-strand breaks (DSBs) comprises multiple steps; sensing and signalling the lesion, mediating the correct type of repair, clearing repair proteins and reforming chromatin. The response involves a diverse set of signalling pathways and repair mechanisms co-ordinated by post-translational modifications (PTMs), phosphorylation, acetylation, ubiquitination, SUMOylation and others.

A major consequence of SUMOylation is the promotion of protein: protein interactions mediated by simple and short hydrophobic SUMO interaction motifs (SIMs) on proximal proteins (Hecker et al., 2006; Psakhye and Jentsch, 2012; Song et al., 2004). In the yeast DNA damage response a SUMO conjugation wave brought about by the interaction of the E3 SUMO ligase Siz2 with DNA and Mre11 results in modification of protein groups thereby promoting SUMO-SIM interactions between members of those groups (Chen et al., 2016; Jentsch and Psakhye, 2013; Psakhye and Jentsch, 2012). In humans, modification by SUMO E3 ligases PIAS1/4 and CBX4 coordinate the repair response, driving the localisation, activity and stability of many signalling and repair proteins, such as RNF168, BRCA1, XRCC4, and Ku70 (Danielsen et al., 2012; Galanty et al., 2009; Hang et al., 2014; Ismail et al., 2012; Lamoliatte et al., 2014; Li et al., 2010; Luo et al., 2012; Morris et al., 2009; Tammsalu et al., 2014; Yin et al., 2012; Yurchenko et al., 2008; Yurchenko et al., 2006). It also fosters key steps such as SUMO-BLM and SUMO-RPA70/RPA1 mediated promotion of RAD51 accumulation (Dou et al., 2010; Eladad et al., 2005; Ouyang et al., 2009; Shima et al., 2013). Many DSB repair factors are SUMOylated, but we currently lack understanding of specific roles for many of these modifications (reviewed in (Garvin and Morris, 2017)).

In the DSB repair response SUMOylation is closely integrated with ubiquitin (Ub) signalling and the Ub-proteasome system (reviewed in (Morris and Garvin, 2017)). This involves SUMO-targeted ubiquitin ligases (STUbLs), which bear tandem SIM motifs and recognize poly-SUMOylated or multi-mono-SUMOylated proteins and target them for ubiquitination and subsequent degradation. Human STUbLs include RNF111/Arkadia (Poulsen et al., 2013) and RING finger 4 (RNF4) (Tatham et al., 2008). Processing of SUMOylated proteins by RNF4 is part of the correct progression of DSB signalling and SUMOylation of MDC1, RIF1 and BRCA1-BARD1 result in their interaction with RNF4 and subsequent degradation after DNA damage (Galanty et al., 2012; Kumar and Cheok, 2017; Kumar et al., 2017; Luo et al., 2012; Vyas et al., 2013; Yin et al., 2012). RNF4 may also regulate RPA residency on ssDNA (Galanty et al., 2012; Yin et al., 2012).

Enzymes with the ability to counter SUMO and Ub modifications have the potential to regulate DNA damage signalling and DNA repair. However, since many SUMOylated factors, and the SUMO machinery itself (Kumar et al., 2017) are eventually processed by STUbLs and degraded by the proteasome, it is also possible that the reversal of SUMO conjugation plays only a minor role in the response. Characterisation of de-ubiquitinating enzymes has shown tremendous diversity and complexity in ubiquitin regulation of the response (Nishi et al., 2014; Uckelmann and Sixma, 2017) but the extent of deSUMOylation enzyme involvement is not known.

Here we establish two mechanisms of DSB repair regulation by the Sentrin Specific Protease 2, SENP2. Firstly we uncover a specific requirement for SENP2 in promoting early DSB signalling by protecting MDC1 from inappropriate SUMOylation and consequent RNF4-VCP processing. We show interaction between SENP2 and MDC1 is released on damage to allow MDC1 SUMOylation required for its clearance. Secondly we reveal that HR repair has a greater need for SUMO conjugates than NHEJ, and thus requires SUMO proteases to contribute to the supply or re-distribution of SUMO. We propose that deSUMOylation is critical to the tuning of both major DSB repair pathways.

## Results

### SENP2 promotes DNA damage signalling and DNA repair

In a prior siRNA screen of SUMO proteases using integrated reporters to measure HR and NHEJ we noted that siRNA to SENP2 resulted in impairment of both repair pathways (Garvin et al., 2013). To address whether SENP2 has a role in DNA repair we compared irradiation (IR) induced γH2AX foci clearance, indicative of DNA repair, and cellular sensitivity to irradiation of wild type (WT) and *SENP2* CRISPR knock out HAP1 cells (*SENP2* KO). *SENP2* KO cells showed both delayed γH2AX foci clearance and greater sensitivity to IR than WT cells (Fig S1A-C).

SENP2 localises to several subcellular compartments, and is enriched at nuclear pores (Chow et al., 2014; Goeres et al., 2011; Hang and Dasso, 2002; Makhnevych et al., 2007; Odeh et al., 2018; Panse et al., 2003; Tan et al., 2015; Zhang et al., 2002). We generated a siRNA-resistant, SENP2^WT^, catalytic mutant (C548A) and a mutant with reduced nuclear pore targeting (NPm - as previously described (Goeres et al., 2011; Odeh et al., 2018) illustrated in S1D). Depletion of SENP2 in HeLa resulted in radio-sensitivity that could be complemented with siRNA-resistant SENP2^WT^ and SENP2^NPm^ but not by SENP2^C548A^ in colony assays (Fig 1A, S1E). Survival in response to Camptothecin (CPT) and Olaparib and measures of both HR and NHEJ repair were also dependent on the catalytic activity of SENP2 (Fig S1F, Fig 1B-C). These data illustrate a need for catalytically competent SENP2 in DNA DSB repair.

**Figure 1.**
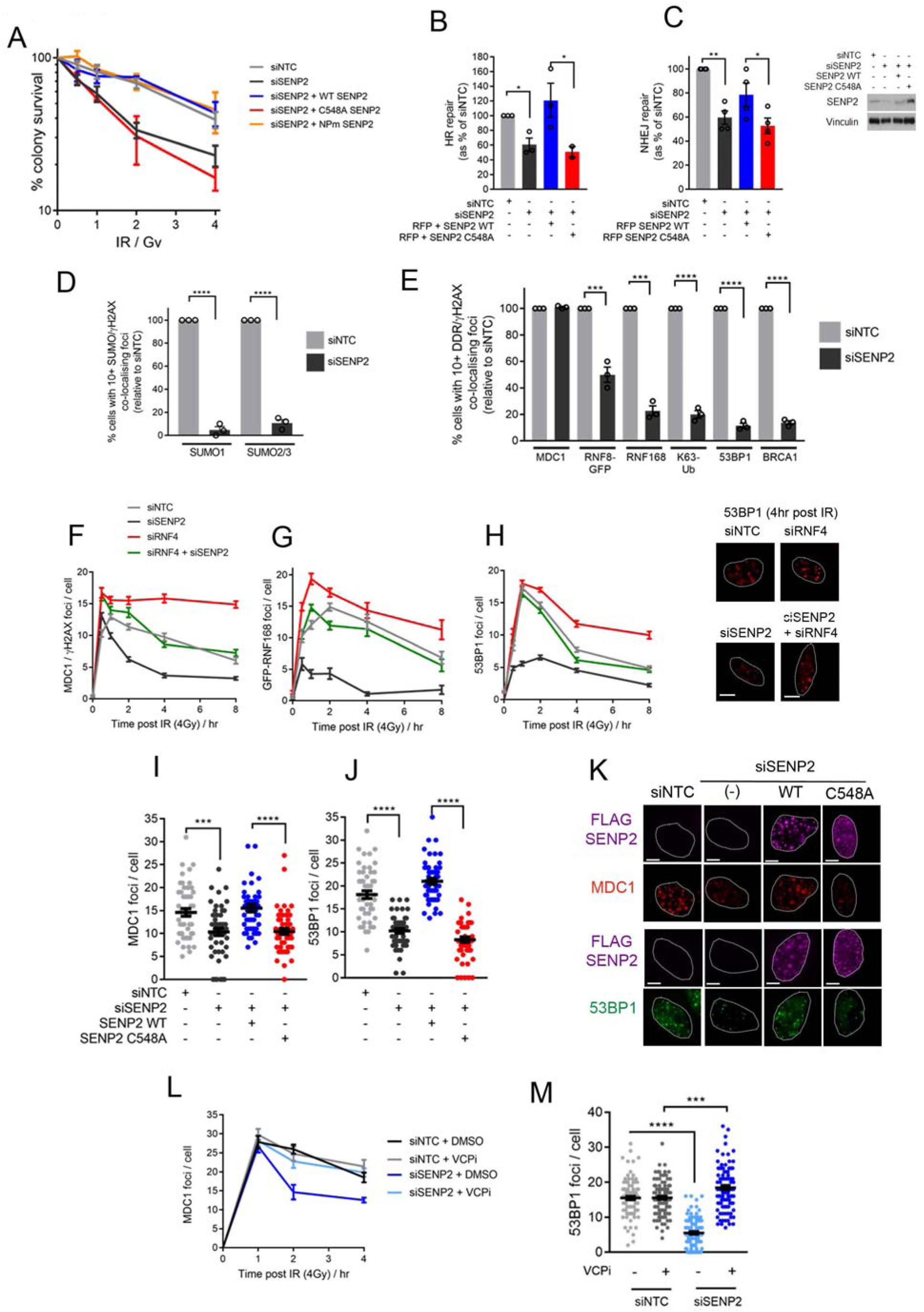
SENP2 promotes DNA damage signalling and DNA repair. **A.** IR colony survival in HeLa treated with siNTC or siSENP2 for 72 hr. Cells were treated concurrently with dox (1μg/mL) to induce siRNA resistant forms of SENP2, n=4. **B-C.** HR (U2OS DR3-GFP) or NHEJ (U2OS-EJ5-GFP) assays using siSENP2 or siNTC treated cells transfected with RFP, *I-SceI* and SENP2^WT^ or SENP2^C548A^. GFP+ cells were normalised to RFP-transfection efficiency. %-repair is given compared to siNTC. Western blot shows SENP2 knockdown efficiency and restoration with siRNA resistant cDNA, n=3. **D-E.** SUMO / γH2AX co-localising foci in HeLa siNTC or siSENP2 cells fixed 1 hr post 5 Gy IR. **E)** as for D with indicated DDR factors, n=3. **F-H.** Time course of MDC1 (n=200), GFP-RNF168 (n=50) or 53BP1 (n=150) foci in HeLa treated with indicated siRNA. Representative images for 53BP1 foci at 4 hr post IR are shown. **I-K.** MDC1 and 53BP1 foci/cell respectively 4 hr post 4 Gy IR in siNTC or siSENP2 HeLa. **K)** representative images related to I), n=100 cells. **L-M.** HeLa (siNTC / siSENP2) irradiated with 4 Gy and 0.5 hr later treated with DMSO / 0.1μM VCPi, CB-5083. Cells were fixed at the indicated times and scored for MDC1 foci. **M)** As for L but 53BP1 foci in cells fixed at 2 hr, n=100 cells.

SUMO1 and SUMO2/3 co-localise with γH2AX foci in response to genotoxic stress such as IR (Galanty et al., 2009; Morris et al., 2009), however following IR we observed less SUMO co-localisation in siSENP2 cells (Fig 1D & S1G). Since a potential cause of SUMO conjugate loss at DSBs is a reduction in the recruitment of proteins on which SUMOylation occurs (Galanty et al., 2009; Morris et al., 2009) we examined cells for DSB repair factor foci. MDC1 is recruited to γH2AX and begins a Ub signalling cascade involving the E3 Ub ligases RNF8/RNF168 to promote the recruitment of the BRCA1-A complex and 53BP1-complex (reviewed in (Panier and Boulton, 2014)). In siSENP2 cells MDC1 co-localization with γH2AX was observed shortly after IR, however RNF8, RNF168, Ub conjugates linked through lysine-63 (K63-Ub), 53BP1 and BRCA1 showed incomplete, or severely reduced, recruitment (Fig 1E). Together these data indicate a role for SENP2 in early DSB signalling.

### RNF4-VCP is responsible for defective DNA damage signalling in SENP2 depleted cells

To determine the signalling breakpoint in SENP2 deficient cells we examined MDC1, GFP-RNF168 and 53BP1 foci kinetics following IR. Depletion of SENP2 severely reduced the accumulation of 53BP1 and RNF168 foci throughout the time course, however MDC1 foci initially formed in siSENP2 and *SENP2-KO* cells and then rapidly became undetectable (Fig 1F & S1H-I). The formation of both MDC1 and 53BP1 foci at later time points, 4 hours after IR, were restored in SENP2^WT^ but not SENP2^C548A^ complemented cells (Fig 1I-K) suggesting deSUMOylase activity is important to the persistence of MDC1 at sites of damage and to the accumulation of 53BP1 foci.

To address which factor(s) are responsible for the rapid clearance of MDC1 in SENP2 deficient cells we first investigated RNF4, whose activity has been implicated in MDC1 turn-over (Galanty et al., 2012; Hendriks et al., 2015; Hendriks and Vertegaal, 2015; Luo et al., 2012; Yin et al., 2012). Co-depletion of RNF4 with SENP2 resulted in foci kinetics of MDC1, RNF168 and 53BP1 similar to that of control-treated cells (Fig 1F-H). The pattern of total SUMO conjugates seen follow IR suggested a similar relationship between SENP2 and RNF4. Control cells exhibited a global increase in high molecular weight SUMO conjugates, particularly for SUMO2/3, after treatment (Fig S2A-B). Whereas in siSENP2 cells, SUMO conjugates were constitutively higher in untreated cells and showed only a slight increase after IR (Fig S2A-B), consistent with the observation of poor DDR protein recruitment and SUMO IRIF formation. Conjugate patterns after siRNF4+siSENP2 co-depletion resembled those seen in siNTC cells (Fig S2A-B) consistent with the near normal DDR foci kinetics observed on co-depletion. Intriguingly loss of the closely related protease, SENP1, did not have a similar impact on SUMO conjugates and depleted cells showed an exaggerated induction of SUMO conjugates following IR (Fig S2C).

RNF4 dependent substrate ubiquitination is frequently followed by processing through VCP (Valosin Containing Protein) hexameric AAA ATPase (Dantuma et al., 2014; Torrecilla et al., 2017). We compared the effects of proteasome (MG132) or VCP inhibition (CB-5083) on MDC1 foci loss after IR. As proteasome inhibition depletes the free Ub pool, in turn causing a failure in Ub signalling in DSB repair (Butler et al., 2012), we also transfected the cells with myc-Ub. MG132 treatment resulted in increased MDC1 foci retention, but in cells expressing additional myc-Ub, foci numbers were reduced, suggesting Ub, rather than the proteasome is critical to MDC1 foci clearance (Fig S2D-E). In contrast, MDC1 foci persistence in the presence of VCP inhibition was unaffected by Ub expression (Fig S2D-E). Moreover in SENP2 depleted cells, the addition of CB-5083 restored near-normal MDC1 foci kinetics and the ability to support downstream 53BP1 foci (Fig 1L-M). Thus RNF4-VCP contributes to the rapid MDC1 foci kinetics in SENP2 deficient cells.

In a further test for potential nuclear pore involvement we examined cells depleted for nuclear pore sub-complex components and known SENP2 interacting proteins; NUP153 and NUP107 (Goeres, 2011 #8625). Reduction in NUP107, had no effect on MDC1 kinetics, and NUP153 depletion modestly increased foci clearance (Fig S2F), confirming no substantial involvement of the nuclear pore in MDC1 kinetics. In contrast when we co-depleted the ligase responsible for MDC1 SUMOylation, PIAS4 (Luo et al., 2012), we found that siPIAS4 (but not siPIAS1), slowed MDC1 foci clearance in siSENP2 cells (Fig S2G). These data consolidate the notion that SUMOylation contributes to the rapid loss of MDC1 foci observed on SENP2 loss.

### MDC1 is a SENP2 substrate and hypo-SUMOylation of MDC1 permits DDR signalling

Lysine 1840 is the main SUMO acceptor site on MDC1 (Fig S3A). To test if MDC1 is a substrate of SENP2 we generated cells expressing myc-MDC1^WT^ or MDC1^K1840R^ and assayed foci kinetics in SENP2 depleted cells. MDC1^WT^ underwent accelerated clearance in siSENP2 cells, as observed for endogenous MDC1. However MDC1^K1840R^ was resistant to the effects of siSENP2, showing the same foci retention in control and siSENP2 cells. Further, expression of this mutant also permitted the formation of downstream 53BP1 foci in siSENP2 treated cells (Fig 2A-B & S3B). Since loss of the main MDC1 SUMOylation site renders damage signalling resistant to the effects of SENP2 repression, these data suggest the impact of SENP2 loss occurs through modification at K1840-MDC1.

**Figure 2.**
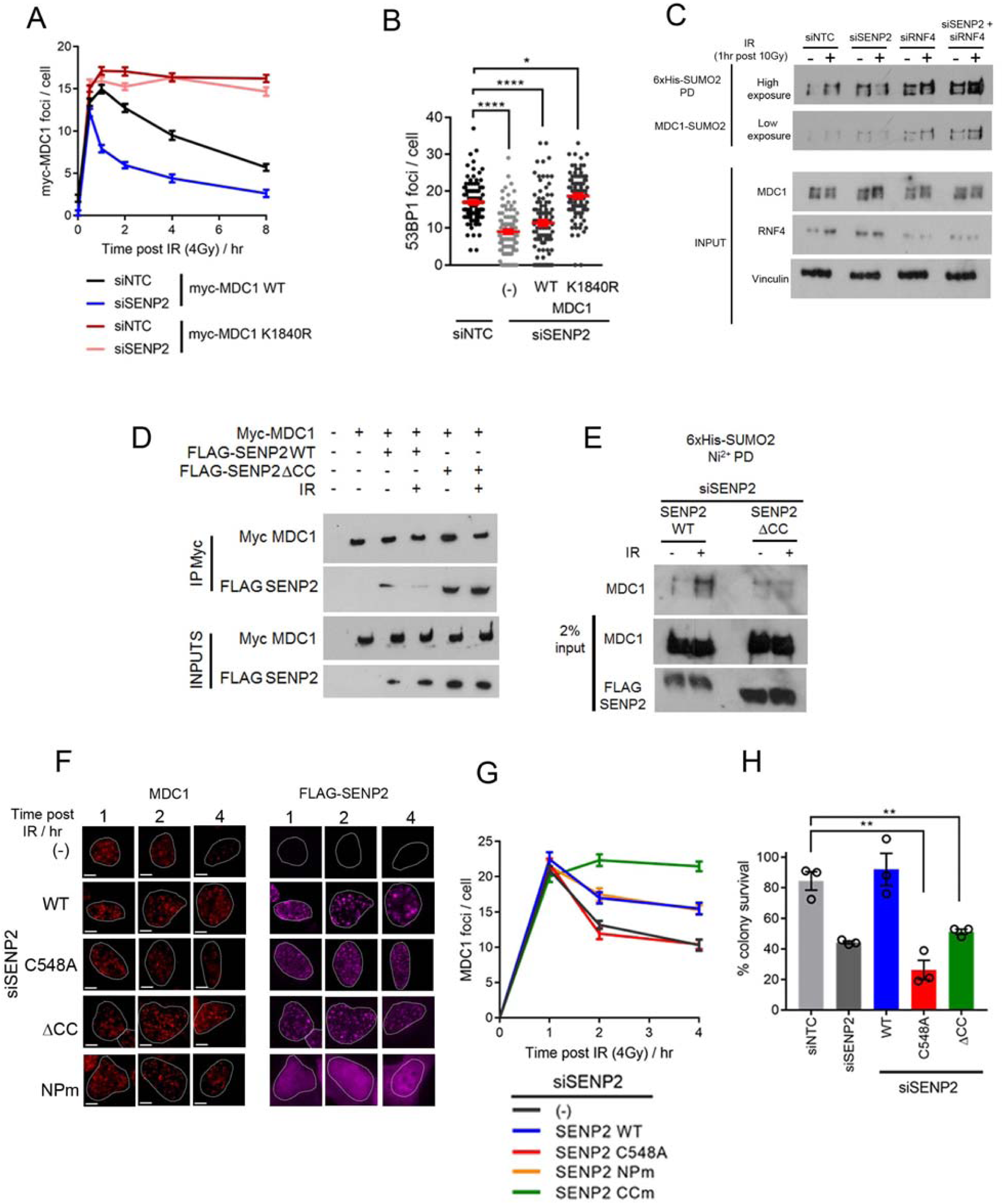
MDC1 is a SENP2 substrate and hypo-SUMOylation of MDC1 permits DDR signalling. **A-B.** HeLa treated with siRNA and induced with dox (72 hr) to express WT or K1840R myc-MDC1. Data shows kinetics of foci/cell for the indicated times post treatment with 4 Gy IR. **B)** As for A but 53BP1 foci at 2 hr. n=100. **C**. HEK293 6x-His-myc SUMO2 treated with indicated siRNA (48 hr), irradiated (10 Gy), lysed 1 hr later and subjected to Ni^2+^ agarose purification, followed by immunoblotting with MDC1 antibodies to determine the relative enrichment in SUMO2 conjugates. PD = Pulldowns. **D**. HEK293 myc-MDC1^WT^ transiently transfected with FLAG-SENP2^WT^ or SENP2ΔCC and treated with dox (72 hr). Cells were irradiated (4 Gy) and lysed 1 hr later followed by immunoprecipitation with myc-agarose. **E**. As for C, but cells were transfected 24 hr post siRNA knockdown with SENP2^WT^ or SENP2ΔCC. **F-G.** HeLa treated with siSENP2. 24 hr later cells were transfected with FLAG-SENP2 for 48 hr, irradiated (4 Gy) and fixed at indicated times. **G)** MDC1 foci/cell were measured in cells co-staining with FLAG-SENP2, n=50. **H**. Colony survival in IR (2 Gy) HeLa treated with siNTC, siSENP2 or siSENP2 plus dox to induce expression of SENP2 mutants, n=3.

We purified His_6_-SUMO1 and His_6_-SUMO2 from untreated and IR-treated cells (harvested 1 hour after IR to capture the MDC1 clearance phase) to test the impact of SENP2 on MDC1-SUMOylation. In untreated siSENP2 or *SENP2-KO* cells we observed an enrichment of MDC1 in SUMO2 conjugates (Fig 2C & S3C-E). Following exposure to IR, cells with SENP2 deficiency exhibited a reduction in SUMOylated MDC1, whereas control cells showed an increase in SUMOylated MDC1 (Fig 2C & S3C-E). In siRNF4+siSENP2 co-depleted cells the IR-dependent reduction of MDC1-SUMO2, seen in siSENP2 cells, was not observed, and instead increased MDC1-SUMO2 was evident as in control cells (Fig 2C & S3C-E). We also confirmed directly that SENP2 could deSUMO2ylate a fragment of MDC1 encompassing K1840 in *vitro* using recombinant SENP2 catalytic domain. (Fig S3F). Next we tested if MDC1 and SENP2 interact and found immunoprecipitated myc-MDC1 co-purified with FLAG-SENP2 in untreated cells, but intriguingly co-precipitation was decreased after IR (Fig 2D). Together these data suggest SENP2 interacts with and restricts MDC1 SUMOylation in untreated cells.

### A conserved coiled-coil region of SENP2 contributes to MDC1 regulation

In a search for regions of SENP2 that may contribute to regulation of MDC1-SUMO, we noted a conserved coiled-coil (CC) domain (Fig S4A-B). We generated a 28 aa deletion mutant, removing the region (ΔCC) and found no changes in protein localisation or activity (Fig S4C-F). However, unlike SENP2^WT^ this mutant retained interaction with MDC1 after exposure to IR (Fig 2D). In complementation assays, SENP2^WT^ permitted increased MDC1 SUMO-2ylation after IR, but cells expressing SENP2ΔCC failed to increase MDC1 SUMOylation (Fig 2E). Moreover cells complemented with SENP2ΔCC failed to clear MDC1 foci and were radiosensitive (Fig 2F-H). These data suggest that dissociation of SENP2 from MDC1 requires the SENP2 CC domain and that dissociation is essential for the IR dependent SUMOylation of MDC1, foci resolution and proper IR repair.

### Requirement for SENP2 can be bypassed by increased K63-Ub signalling

We observe an initial association of MDC1 at DSBs in siSENP2 cells (Fig 1F & S1H-I), leading to the question of what element of the DDR is effected by rapid loss of MDC1 from damage sites. Intriguingly a similar impact is seen on DSB signalling when MDC1 turn-over is increased, but steady-state foci are only slightly altered, following loss of the DUB Ataxin-3 (ATXN3) (Pfeiffer et al., 2017). We note that SENP2 and ATXN3 both contribute to the longevity of MDC1 foci and colony survival in response to IR (Fig S5A-B), so that together these observations suggest that MDC1 residency, or quality of MDC1 at sites of damage promotes downstream signalling. MDC1 is involved in two positive feedback loops that may require its prolonged association. It contributes to signal amplification of γH2AX around DNA break sites with MRN and ATM (Chapman and Jackson, 2008; Savic et al., 2009; Stucki et al., 2005) and to the amplification of K63-Ub linkages on Histone H1 and L3MBTL2 (Nowsheen et al., 2018; Thorslund et al., 2015) with RNF8 and, downstream, RNF168 (reviewed in (Panier and Boulton, 2014)). Since we observed no loss of γH2AX foci in SENP2 deficient cells (Fig S1A, S1H) we tested whether insufficient K63-Ub generation contributes to DDR signal failure by manipulating the K63-Ub machinery. Over-expression of RNF8, which catalyses the initial K63-Ub contribution (Lok et al., 2011; Thorslund et al., 2015) and the depletion of the K63-Ub specific ubiquitin protease, BRCC36 (depletion of which increases K63-Ub at sites of damage (Shao et al., 2009)) were capable of restoring 53BP1 foci in siSENP2 cells (Fig S5C-E). These data suggest normal turn-over kinetics of MDC1 at damage sites is needed for sufficient Ub conjugate generation.

### RNF4-VCP is responsible for the IR-sensitivity of SENP2 depleted cells

Prompted by our findings that RNF4 is responsible for rapid MDC1 foci kinetics in siSENP2 depleted cells we next assessed if RNF4 contributes to their radiosensitivity. Depletion of RNF4 or SENP2 individually increased cell sensitivity to IR, but co-depletion resulted in IR resistance similar to siNTC cells (Fig 3A). Expression of RNF4^WT^ restored resistance to RNF4 depleted cells, however, critically, on a siSENP2 + siRNF4 background re-introduction of RNF4^WT^ resulted in IR sensitivity (Fig 3B) demonstrating the toxicity of RNF4 in siSENP2 cells. Complementation with RNF4 proteins that reduce interaction with Ub loaded E2 conjugating enzyme, prevent RNF4 dimerization or interaction with SUMO (Kung et al., 2014; Plechanovova et al., 2012; Rojas-Fernandez et al., 2014) allowed survival on siRNF4 + siSENP2 backgrounds, but not cells treated with siRNF4 alone (Fig 3B). We confirmed the corollary of these findings; that SENP2 protease activity contributes to the toxicity of IR in siRNF4 cells (Fig 3C). Moreover VCP inhibition restored IR-resistance to siSENP2 cells (Fig 3D). Thus the SUMO-targeting and Ub ligase function of RNF4 and VCP activity contributes to the IR sensitivity of SENP2 depleted cells. Amongst the SENP family of SUMO proteases SENP2 is alone in contributing significantly to the lethality of IR in RNF4 depleted cells (Fig S5F). Together our data reveal a strong reciprocal relationship between RNF4 and SENP2 in the cellular response to IR, consistent with their opposing influences on MDC1 in DSB damage signalling.

**Figure 3.**
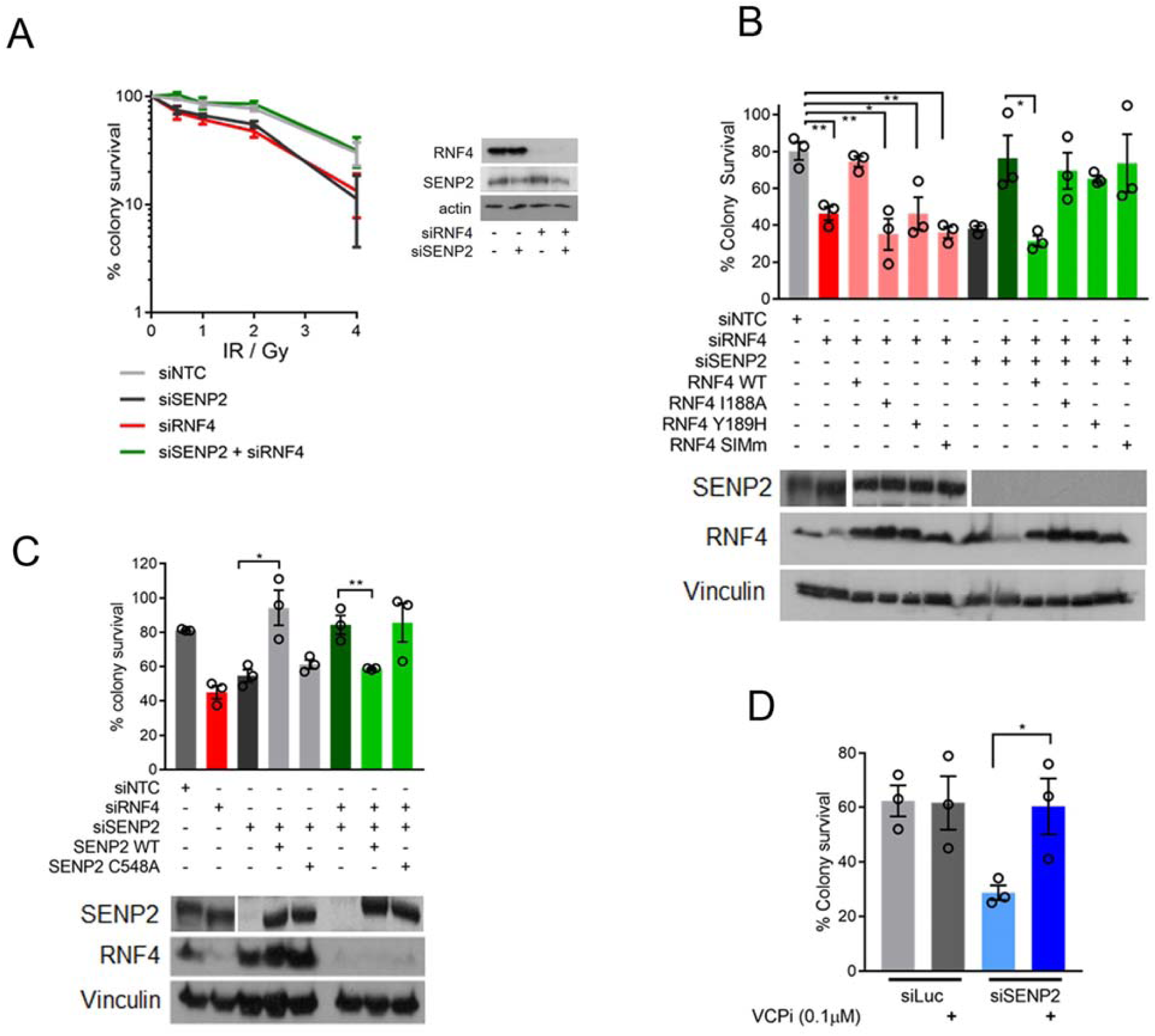
RNF4-VCP is responsible for the IR-sensitivity of SENP2 depleted cells. **A**. IR colony survival HeLa treated with indicated siRNA. Right panel western blot of siRNA depletions. **B**. Colony survival in HeLa RNF4 treated with indicated siRNA and dox to induce expression of RNF4 and its mutants for 72 hr prior to 2 Gy IR, n=3. RNF4 antibody will also detect exogenous protein. **C**. As for B, but using HeLa SENP2, n=3. SENP2 antibody will also detect exogenous protein. **D**. Colony survival of HeLa treated with siRNAs shown and 4 Gy IR. Thirty minutes post irradiation cells were treated with DMSO, VCP inhibitor CB-5083 (0.1μM), n=3.

### SENP2 is not relevant to S-phase clearance of MDC1

We expected to see a role for SENP2 in regulating MDC1 at repair foci throughout the cell cycle. However, when we labelled cells with a nucleotide analogue to differentiate S-phase cells we found that SENP2 depletion had no influence on MDC1 in S-phase marked cells (Fig 4A-C). Expression of the MDC1^K1840R^ SUMO-site mutant results in cellular IR sensitivity, due to a failure of the mutant to clear from sites of DNA damage (Luo et al., 2012). We confirmed these data (Fig 4D-E), but also challenged cells with CPT and Olaparib, agents that require HR repair for resistance, and found MDC1^K1840R^ did not increase sensitivity to these agents (Fig 4F-G). Moreover the MDC1^K1840R^ mutant had no negative impact on RAD51 foci formation in S-phase cells (Fig 4H). While MDC1^K1840R^ expression increased 53BP1 foci numbers in EdU-cells, as previously reported (Luo et al., 2012), it did not alter 53BP1 foci number in EdU+ cells (Fig 4I). Moreover unlike the response to IR, co-depletion of RNF4 and SENP2 did not improve survival of cells challenged by CPT or Olaparib and did not substantially improve HR reporter activity nor improve RAD51 foci accumulations over single depletions (Fig 4J-N). We conclude S-phase cells turn-over MDC1 from broken DNA ends independently of its major SUMO-acceptor site and of SENP2, suggesting the role of SENP2 in HR repair occurs in another pathway.

**Figure 4.**
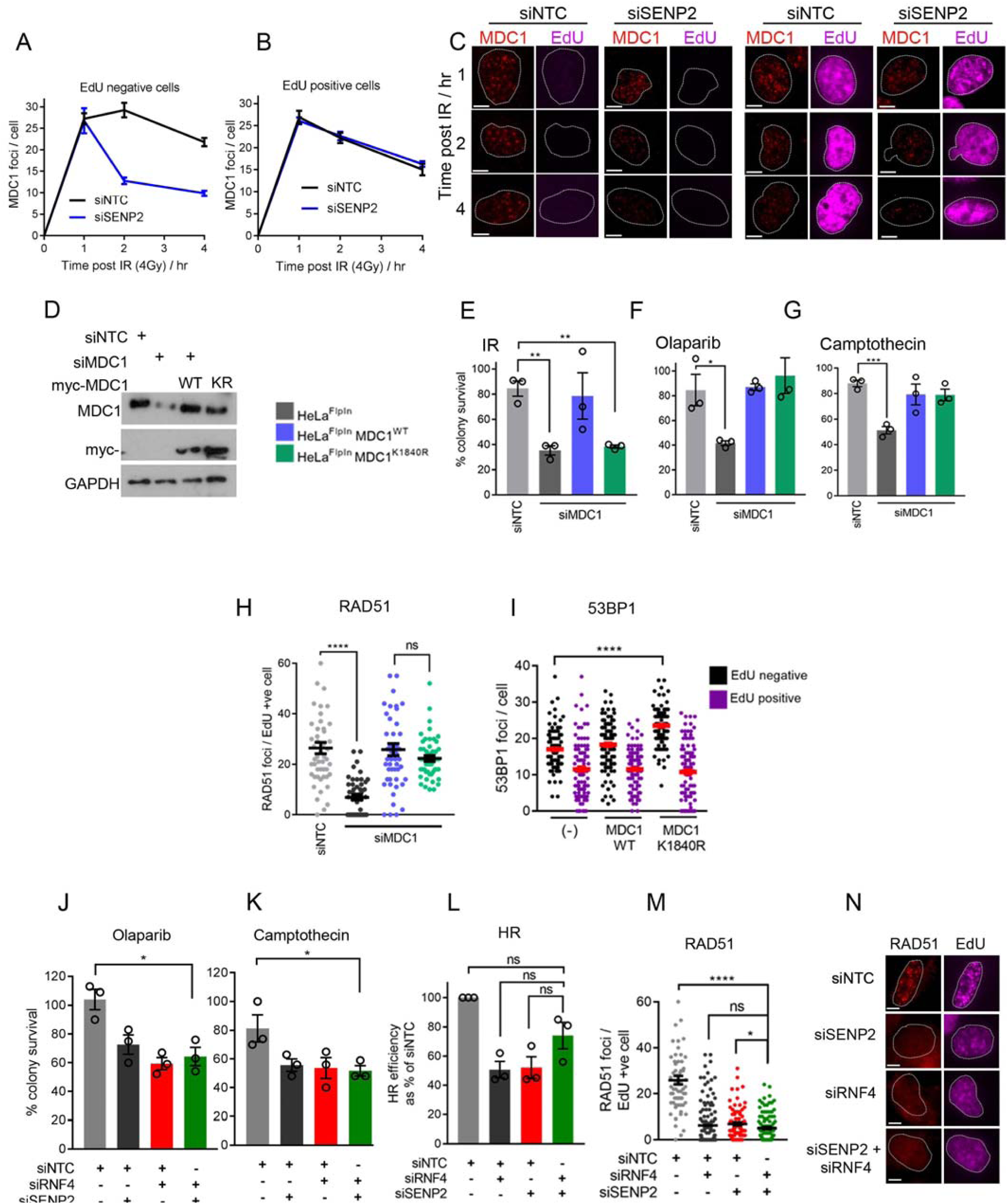
SENP2 is not relevant to S-phase clearance of MDC1. **A-B.** HeLa treated with siNTC or siSENP2 for 72 hr, pulsed with 10μM EdU 1 hr prior to IR (4 Gy). Cells were fixed at indicated times and subjected to Click-It labelling with 647 nm azide to detect EdU incorporation into nascent chromatin. MDC1 foci/cell in (A) EdU negative and (B) EdU positive (S phase) cells. 50 cells were scored per condition from a total of 3 experiments. **C**. Representative images relating to (A-B). **D**. Western blot showing MDC1 knockdown and expression in HeLa myc-MDC1 **E-G.** HeLa myc-MDC1^WT^ and K1840R cells siRNA depleted for endogenous MDC1 and treated with dox to induce MDC1. After 72 hr cells were treated with (E) 2 Gy IR, (F) 10μM Olaparib or (G) 2.5 μM CPT (2 hr) and subjected to colony survival analysis, n=3. **H**. HeLa treated as for (A), but stained with RAD51, n=100 cells. **I**. HeLa treated with dox to induce expression of myc-MDC1^WT^ or K1840R for 72 hr. 1 hr prior to IR (4 Gy) cells were pulsed with EdU and fixed at 2 hr later. Cells (100 from a total of 3 experiments) were scored for 53BP1 foci/cell in EdU -/+ cells. **J-K.** Colony survival in HeLa treated with siRNA and drug as for (F-G), n=3. **L**. U2OS-DR3 homologous recombination reporter cells treated with siRNA for 24 hr prior to transfection with *i-Sce-I* nuclease and RFP (to control for transfection efficiency) for a further 48 hr. The % RFP/GFP positive cells relative to siNTC is shown for 3 experiments. **M**. RAD51 foci in HeLa treated as for (H), n=100. **N**. Images relating to (M).

### Homologous recombination is highly sensitive to the supply of SUMO

Since we observed increased high molecular weight SUMO-conjugates in untreated cells depleted of SENP2 (Fig S2A) we speculated that SENP2 loss may disable HR through reduced availability of SUMO for conjugation. We over-expressed conjugation proficient and deficient SUMO in siSENP2 cells and examined survival in response to IR, CPT or Olaparib. We also assessed MDC1 foci 4 hours after IR and RAD51 foci in S-phase cells. SUMO expression had no influence on IR-resistance nor MDC1 foci (Fig 5A-C) but conjugation competent SUMO isoforms, particularly SUMO2, improved CPT and Olaparib resistance and restored RAD51 foci in SENP2 depleted cells (Fig 5D-F). Intriguingly SENP2 depletion had no impact on RPA foci accrual suggesting a role for SENP2 in RAD51 loading but not DNA end resection (Fig S5G). To test the hypothesis that differential requirements for SUMO availability exist between different repair mechanisms we performed a partial depletion of SUMO2/3 (Fig 5G-H). Remarkably, partial SUMO2/3 depletion resulted in CPT and Olaparib, but not IR, sensitivity and impaired HR but not NHEJ in integrated repair assays (Fig 5I-J), indicating that the HR-pathway is more sensitive to SUMO availability.

**Figure 5.**
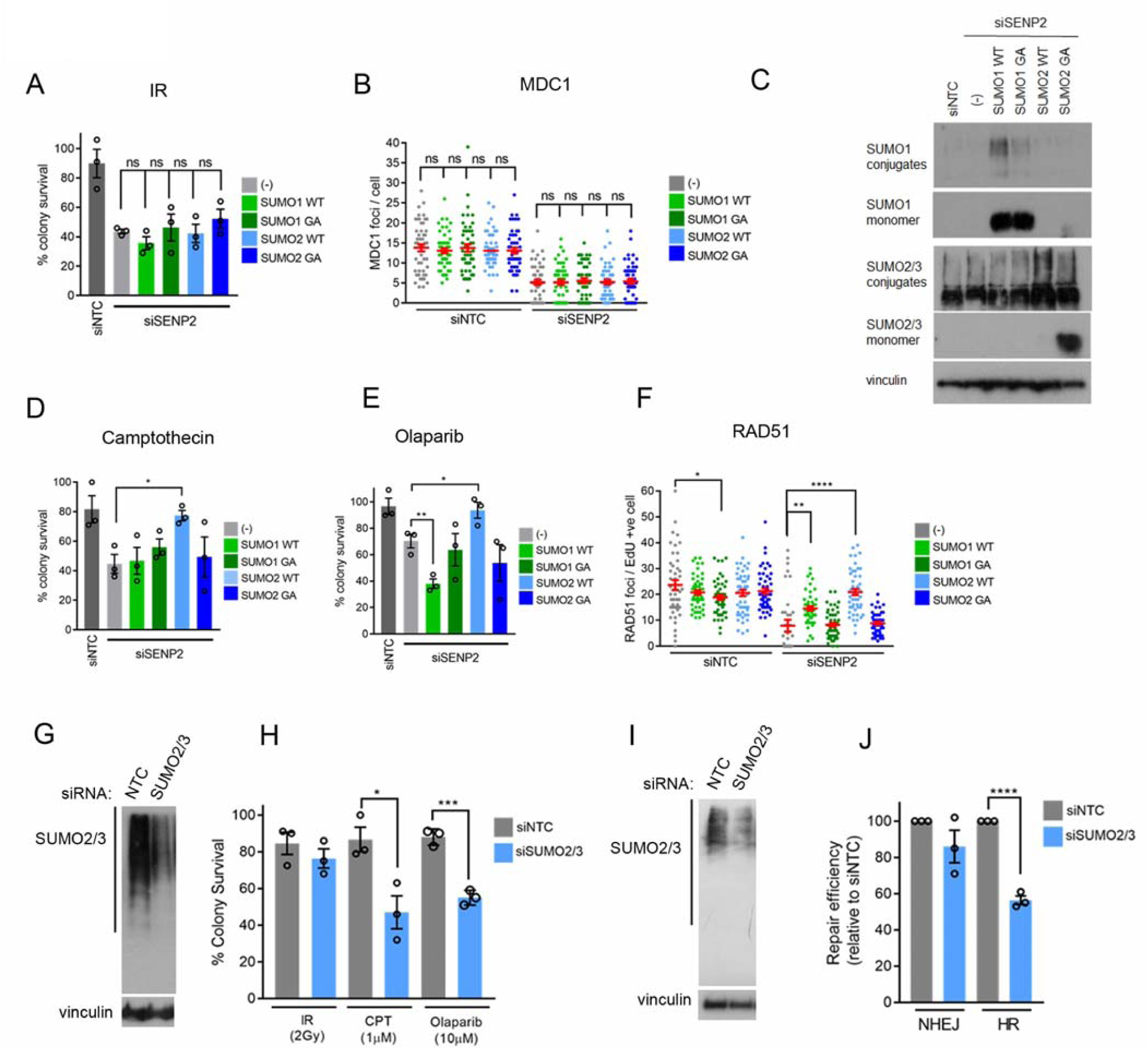
HR is sensitive to the supply of SUMO. **A**. Colony survival after 2 Gy IR in HeLa 6xHis-myc SUMO in siNTC / siSENP2 depleted cells. GA indicates di-glycine → alanine mutants in SUMO isoforms that prevent conjugation, n=3. **B**. HeLa treated with siRNA for 24 hr before transfection with myc-SUMO, cells were treated with 4 Gy IR 48 hr later and immunostained for MDC1 in myc-SUMO expressing cells, n=100. Western blot of SUMO conjugates relating to (A-B). **D-E**. As for (**A**) but using (**D**) 1 μM CPT or (E) 10 μM Olaparib for 2 hr before plating for colony survival, n=3. **F**. As for (B), but 1 hr prior to fixation cells were incubated with 10μM EdU to label replicating cells and stained for RAD51. **G-H.** Western blot showing partial depletion of SUMO2/3 conjugates in HeLa. **H)** Colony survival in HeLa depleted with siNTC or siSUMO2/3 followed by treatment with 2 Gy IR, 1 μM CPT or 10 μM Olaparib. **I-J**. Western blot showing SUMO2/3 knockdown. **J)** U2OS HR and NHEJ reporters treated with siNTC or siSUMO2/3 and transfected with *i-Sce-I* and RFP for 72 hr. HR and NHEJ efficiency was set at 100% for siNTC.

### High levels of SENP2 disrupt DSB repair

The *SENP2* gene maps to chromosome 3q26-29, a region commonly amplified in epithelial cancers of the lung, ovaries, oesophagus and head and neck (Cancer Genome Atlas, 2015; Qian and Massion, 2008) (Fig S6A-B). In lung cancer high SENP2 mRNA levels correlate both with copy number and reduced patient survival (Fig S6C-D). Since our data shows a critical role for SENP2 in DSB repair we explored whether increased SENP2 expression alters repair. Elevation of SENP2 expression resulted in increased 53BP1 and MDC1 dependent resistance to IR (Fig 6A-B) and was accompanied by persistent MDC1 foci at 2 hours after IR (Fig 6C-D). With the exception of SENP6 no other SENP expression slowed MDC1 clearance (Fig S6E-F). High expression of SENP2 also induced a 2.5 fold increase in NHEJ measured from an integrated substrate (Fig 6E-F). Thus increased SENP2 results in slower MDC1 clearance correlating with increased IR resistance and improved NHEJ.

**Figure 6.**
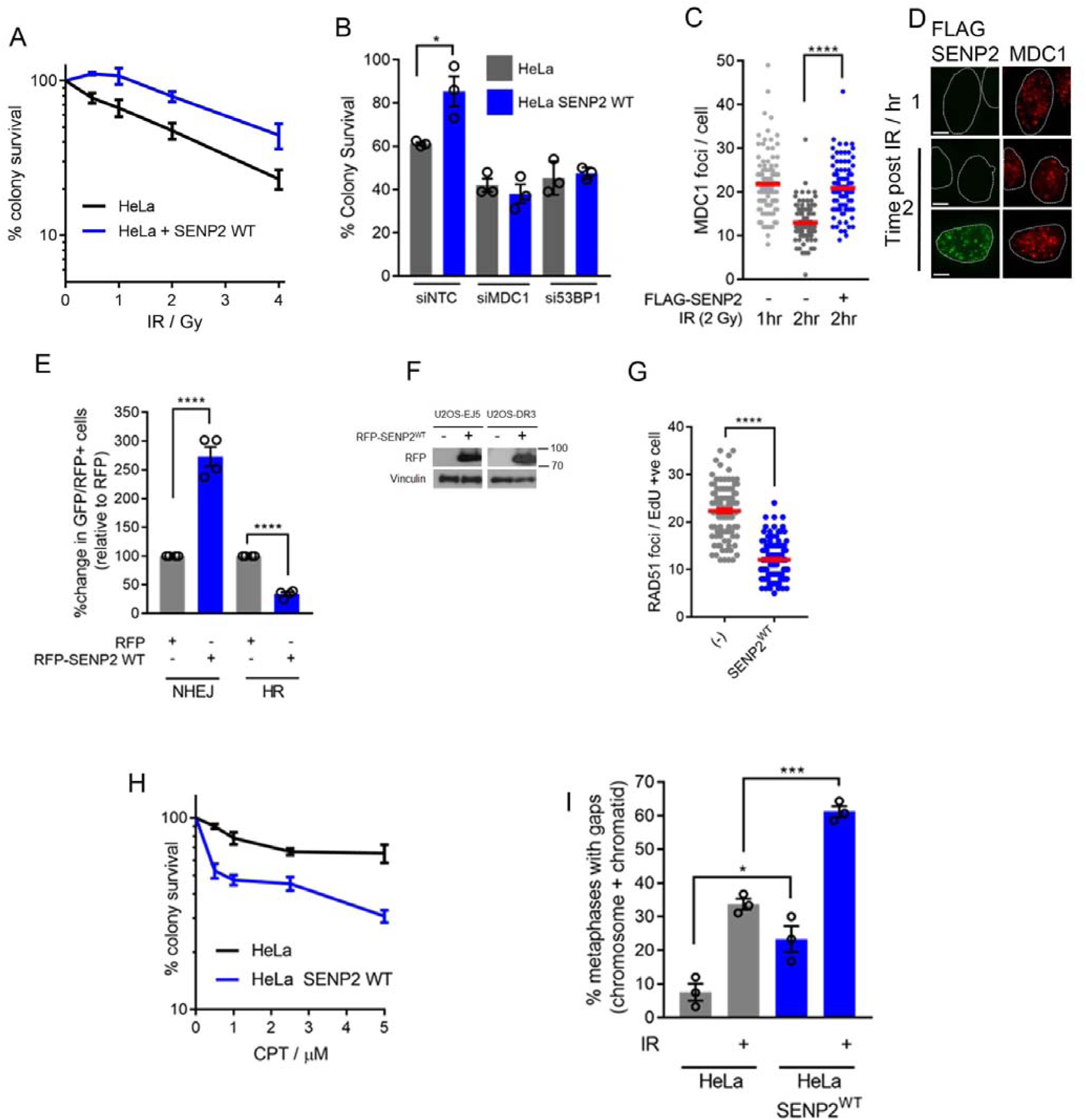
SENP2 over-expression disrupts responses to DSBs. **A-B.** HeLa SENP2^WT^ -/+ dox for 72 hr were treated with indicated dose of IR and subjected to colony survival analysis. (B) Colony assay performed as for (A) but with siRNA transfection concurrent with Dox addition. IR = 2 Gy, n=3. **C-D.** HeLa transfected with SENP2^WT^ for 48 hr prior to 4 Gy IR and fixation 2 hr later. MDC1 foci / cell were scored in FLAG-SENP2 positive cells n=100. **D**) representative images of C. **E-F**. HR and NHEJ U2OS reporters expressing RFP or RFP-SENP2 and *I-Sce1* GFP-positive cells were normalised to RFP-transfection efficiency. %-repair is given compared to NTC. **F**) Western blot showing expression of RFP-SENP2, n=4. **G**. for (**C**) except cells were treated with EdU to label replicating cells. EdU positive cells were scored for the number of RAD51 foci / cell. **H**. As for (**A**) but using 2 hr treatment of 2.5 μM CPT prior to plating. **I**. HeLa SENP2^WT^ or HeLa treated with dox for 72 hr prior to IR 2 Gy, 18 hr later cells were treated with colcimid for 6 hr and processed for metaphase spread analysis. Data shows % metaphases with chromosome/chromatid gaps from 3 experiments.

Increased expression of SENP2 reduces high molecular weight SUMO conjugates (Fig S4E-F) leading us to speculate whether persistent removal of SUMO may also influence HR. High SENP2 expression resulted in reduced HR reporter product, reduced RAD51 foci and reduced CPT resistance (Fig 6G-H). Examination of chromosome aberrations in cells acutely over-expressing SENP2 showed increased chromosomal gaps suggesting an overall reduced repair ability despite improved resistance (Fig 6I).

The 3q amplification carries two more genes involved in DNA repair; the Ub ligase RNF168, and the de-ubiquitinating enzyme USP13 (Fig 7A), which contribute to DNA damage signalling (Chroma et al., 2017; Doil et al., 2009; Li et al., 2017; Nishi et al., 2014; Stewart et al., 2009). We compared the influence of high expression of each repair gene in Hela^FlpIn^ stable doxycycline inducible cells. Of the three genes, SENP2 had the greatest influence on IR-resistance, while increased SENP2 and USP13 reduced CPT resistance (Fig 7B-D)

**Figure 7.**
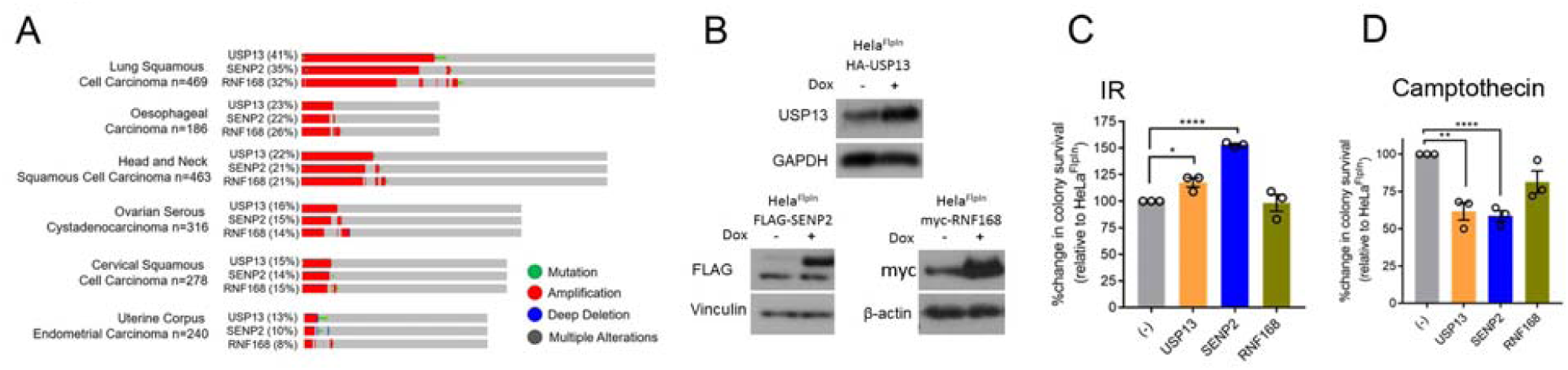
SENP2 is part of a DSB-repair disruptive amplicon on 3q. **A.** Oncoprints adapted from Cbioportal TCGA datasets (August 2018) for USP13, SENP2 and RNF168 genomic amplification (red) in indicated cancer types. Values in parenthesis indicate % of samples with amplification. **B** Western blot of USP13, RNF168 and FLAG-SENP2 expression. **C-D** Colony survival IR (2 Gy) or CPT (1 μM 2 hr) in HeLa over-expressing USP13, SENP2 or RNF168 n=3.

## Discussion

The sequential action of PTMs is essential for the proper cellular response to DSBs. While cross-talk between SUMOylation and ubiquitination is important for the integration of signalling cues for the response, the extent of a role for deSUMOylation was poorly defined. Here we provide evidence that deSUMOylation is required prior to the onset of DSB signalling to govern correct PTM timing following damage. Mechanistically we identify two distinct pathways in which deSUMOylation is required (Fig S7).

We show that interaction between MDC1 and SENP2 in untreated cells is associated with MDC1-hypoSUMOylation and with Ub signalling in the damage response. In the absence of SENP2, PIAS4-mediated SUMOylation facilitates rapid RNF4-VCP-mediated MDC1 turn-over and a failure of down-stream signalling. MDC1 SUMOylation and RNF4-processing is induced on chromatin (Luo et al., 2012) so one question arising from our study is why constitutive interaction with a SUMO protease is needed? MDC1-SUMO is detected in untreated cells (Galanty et al., 2012; Hendriks et al., 2015; Hendriks and Vertegaal, 2015; Luo et al., 2012; Vyas et al., 2013; Yin et al., 2012), suggesting a constitutive propensity to modification, even in the presence of SENP2. Thus interaction with a SUMO protease may be needed to prevent the accumulation of a heavily modified protein capable of driving its own removal.

We show that a novel, conserved coiled-coil region (aa203-228) is needed to release MDC1-SENP2 interactions following IR and to allow subsequent MDC1 SUMOylation needed for its eventual clearance from damaged chromatin. The SENP family of SUMO proteases contain relatively few functionally annotated domains outside of the C-terminal catalytic regions (Mukhopadhyay and Dasso, 2007). How the coiled-coil allows IR-regulated dissociation remains to be discovered. Its conservation in evolution as far as chicken and zebrafish (Fig S4B) suggests an important role for the motif.

Surprisingly we find that the role of SENP2 in S phase has no relationship with the MDC1-processing pathway. Instead measures of HR, repressed in SENP2-deficient cells, are rescued by the expression of exogenous SUMO2/3. SUMO conjugation is required for both main pathways of DSB repair so that a total loss of SUMO availably/conjugation restricts both mechanisms (Galanty et al., 2009; Morris et al., 2009), what is striking in our findings is evidence for a level of SUMO availability at which NHEJ can function but HR cannot. The degree to which each repair process captures available SUMO is not known. The differential requirement may reflect the greater number of SUMOylated factors in HR over NHEJ, a greater need for group-modification in HR, or a greater need for the promotion of particular interactions, for example between BLM, RPA and RAD51 (Bologna et al., 2015; Dou et al., 2010; Eladad et al., 2005; Galanty et al., 2012; Garvin and Morris, 2017; Hendriks and Vertegaal, 2016; Ouyang et al., 2009). Alternatively, cells in S-phase may have greater need for available SUMO in replicative processes, reducing availability to HR.

In the HeLa^FlpIn^ cells used in the current study we detected no free SUMO pool and no accumulation of exogenous SUMO2 in a free SUMO2/3 pool, suggesting the increase in SUMO2 availability was immediately captured within conjugates. In some cell types the vast majority of SUMO exists in conjugates, for example shifting from 93% of SUMO2/3 in conjugates to 96% and 98% on MG132 and heat shock, respectively in HEK293T cells (Hendriks et al., 2018). In these contexts induced SUMO conjugation in stress responses may be reliant on SUMO synthesis and recycling from SUMOylated proteins.

We show that acute, high level expression of SENP2 results in increased NHEJ correlating with extended MDC1 foci longevity and 53BP1-dependent IR-resistance, consistent with extended defence of MDC1-SUMO. SENP2 over-expression also reduces global SUMO-conjugates, and we speculate strips SUMO from HR-proteins during the damage response. In both pathways SENP2 levels dramatically influence repair outcomes.

Many cancers have altered SUMOylation (Seeler and Dejean, 2017) and SENP2 transcription can be upregulated by NF-κB (Lee et al., 2011) activation of which is a hallmark of cancer development. SENP2 is one of several genes on the amplified region of chromosome 3q with the capacity to influence survival to DNA-damaging therapeutics. Evidence of SUMO-pathway addiction, has been found in Myc and Ras driven cancers (Kessler et al., 2012; Yu et al., 2015) while those with low SUMOylation may be sensitive to further targeting of the SUMO system (Licciardello et al., 2015). Differential needs for SUMO conjugation and de-conjugation therefore could expose tumour-specific vulnerabilities. SUMO E1 and E2 inhibitors have been described (He et al., 2017; Kumar et al., 2016; Lu et al., 2010) and our data implies that partial SUMO-conjugation inhibition could disable HR, but not NHEJ, increasing the mutation load relevant to immune blockade and sensitivity to HR-directed therapies. Moreover our data suggest that development of SENP2 inhibitors, beyond currently available tool compounds (Kumar et al., 2014; Madu et al., 2013), could have utility in the treatment of certain common 3q-amplified tumours while sparing normal tissue. Aerodigestive-track cancers often receive post-operative radiotherapy so further investigation into the potential of targeting SENP2 in the context of chromosome 3q amplification is warranted.

In summary the need for the SUMO protease, SENP2, in aspects of mammalian DSB repair presented here reveal unexpected requirements for SUMO deconjugation, and its regulation, in the DNA damage response and place the need for the activity largely in undamaged cells before the stress of DSBs occurs. We find deSUMOylation by SENP2, prevents engagement of RNF4-VCP with MDC1, restricting an ‘over-before-it-has begun’ repair response and promotes SUMO supply, critical to the completion of HR, while increased SENP2 expression dramatically dysregulates DSB repair mechanisms.

## Materials and Methods

### Colony survival assays

Cells were plated at 2.5 x 10^5^ cells/ml in a 24 well plate. For siRNA transfections, cells were transfected 24 hr post plating for an additional 48 hr. For over-expression cells were treated with doxycycline (1μg / mL) for 72 hr. Cells were treated with the indicated dose of DNA damaging agent prior to plating at limiting dilution in 6 well plates to form colonies and grown on for 10 days (3 wells / technical repeat). Colonies were stained 0.5% crystal violet (BDH Chemicals) in 50% methanol and counted. Colony survival was calculated as the % change in colonies versus untreated matched controls. Graphs shown are combined data from a minimum of 3 independent experiments and error bars show SEM.

### Transfections

Small interfering RNA (siRNA) transfections (10nM) were performed using Dharmafect1 (Dharmacon) and DNA plasmids using FuGENE 6 (3 µl:1 µg FuGENE: DNA) (Promega) following the manufacturer’s protocols. SMARTPools were from Dharmacon and individual sequences were from Sigma. See table 3 for siRNA sequences. Cells were grown for 48-72 hr post-transfection before treatment and harvesting.

### Drug treatments

Irradiation was performed with a Gamma-cell 1000 Elite (Cs^137^) radiation source. The following chemicals were used, CB-5083 / VCPi (0.1 μM) (Selleck chemicals), Camptothecin (1 μM) (Sigma), Olaparib (10 μM) (Selleck chemicals), EdU (10 μM) (Life Technologies), MG132 (10 μM) (Sigma), Colcimid (Sigma).

### NHEJ and HR assays

U2OS-DR3-GFP (gene conversion), and U2OS-EJ5-GFP (Non-homologous end-joining) were a generous gift from Jeremy Stark (City of Hope, Duarte USA). U20S reporter cell lines were simultaneously co-transfected with siRNA using Dharmafect1 (Dharmacon) and DNA (RFP, or Flag-SENP2 and *I-Sce1* endonuclease expression constructs) using FuGene6 (Promega) respectively. After 16 hr the media was replaced and cells were grown for a further 48 hr before fixation in 2% PFA. RFP and GFP double positive cells were scored by FACS analysis using a CyAn flow cytometer and a minimum of 10000 cells counted. Data was analysed using Summit 4.3 software. Each individual experiment contained 3 technical repeats and normalized to siRNA controls or to WT-complemented cells. Graphs shown are combined data from a minimum of 3 independent experiments and error bars show standard error.

## Immunofluorescence

Cells were plated on 13 mm circular glass coverslips at a density of 5 x 10^4^ cells/ml, treated as required. For RPA, and RAD51 staining cells were pre-extracted in CSK buffer (100 mM sodium chloride, 300 mM sucrose, 3 magnesium chloride, 10 mM PIPES pH 6.8) for 1 minute at room temperature, For all other staining’s cells were first fixed in 4% PFA and permeabilised with 0.5% Triton X100 in PBS. After blocking in 10% FCS, cells were incubated with primary antibody for 1 hr (unless otherwise stated) and with secondary AlexaFluor antibodies for 1 hr. The DNA was stained using Hoechst at 1:20,000. In some images the DNA stain has been drawn around (but not shown) to illustrate the location of the nucleus.

RAD51 foci: Cells were labelled with 10 µM EdU 1 hr prior to IR using a Gamma-cell 1000 Elite irradiator (caesium-137 source). At 4 hr post-IR cells were washed briefly in CSK buffer (100 mM sodium chloride, 300 mM sucrose, 3 mM magnesium chloride, 10 mM PIPES pH 6.8) before fixation with 4 % Paraformaldehyde for 10 min. For IF staining cells were permeabilised with 0.2% TritonX100 in PBS for 10 min before blocking in 10 % FBS in PBS. EdU was visualised by Click-iT^®^ chemistry according to the manufacturer’s protocols (Life Technologies) with Alexa-647-azide. Cells were incubated with primary antibody overnight, washed three times in PBS and incubated with secondary AlexaFluor antibodies for 1 hr.

With the exception of Figure 1G-H all immunofluorescent staining was imaged using the Leica DM6000B microscope using a HBO lamp with 100W mercury short arc UV bulb light source and four filter cubes, A4, L5, N3 and Y5 to produce excitations at wavelengths 360 488, 555 and 647 nm respectively. Images were captured at each wavelength sequentially using the Plan Apochromat HCX 100x/1.4 Oil objective at a resolution of 1392×1040 pixels. Detection of SUMO IRIF was performed according to (Morris 2009).

### Cloning

**SENP2** was cloned with an N terminal FLAG tag into the *KpnI* and *EcoRV* sites in pCDNA5/FRT/TO vector (Invitrogen). Synonymous mutations were made in the SENP2 cDNA to generate siRNA resistance (see table 4). SENP2 cDNA was also cloned into pCDNA3.1 mRFP vector using *ClaI*. All site directed mutagenesis was performed using Pfu polymerase (Promega) and mutations were confirmed by Sanger sequencing (Source Biosciences Nottingham). To generate a nuclear pore binding mutant of SENP2 we truncated amino acids 1-65 and mutated the SENP2 NES to prevent nuclear export. The coiled coil deletion mutant was generated using the megaprimer method with primers that flank the deleted region and external primers to generate the megaprimer. The PCR product was then used for site directed mutagenesis. **MDC1**, the longest isoform of human MDC1 (NM_014641.2) was used to generate synthetic MDC1 cDNA that was extensively codon optimised by GenScript to remove repetitive DNA sequences to enable gene synthesis. The optimised cDNA has an N terminal myc tag, synonymous mutations to enable resistance to two siRNA targeting Exon 11 and multiple silent mutations that disrupt restriction enzyme recognition sites. The myc-MDC1 cDNA was cloned into *AflII* and *BamHI* sites in pCDNA5/FRT/TO. The K1840R mutation was made by GenScript. To generate the MDC1 fragments for in vitro SUMOylation / deSUMOylation, WT and K1840R MDC1 were and cloned into pCA528 containing a His-SUMO n terminal tag using BsaI and BamHI sites. **RNF4,** human RNF4 (NM_002938.4) cDNA was synthesised by GenScript to contain resistance to two siRNA sequences, an N terminal HA tag, and cloned into pCDNA5/FRT/TO HindIII and BamHI sites. Site directed mutagenesis was used to generate the RNF4 mutants. The SIM mutant of RNF4 was generated by SDM of SIM2 and SIM3 followed by the megaprimer method using a forward primer that contained mutations in SIM1 and a reverse primer that contained mutations in SIM4. **RNF168** was cloned from pEGFP-RNF168 (a kind gift of Grant Stewart, University of Birmingham). The two BamHI sites were silenced with synonymous mutations by site directed mutagenesis, and the resulting cDNA was sub-cloned into pCDNA5/FRT/TO using BamHI-XhoI sites. **SUMO1 and SUMO2 (**NM_003352.4, NM_006937.3) cDNA (both in their processed forms) were cloned into pCDNA5/FRT/TO with an N terminal 6x Histidine - myc tag. GA mutations that prevent SUMO conjugation were generated by incorporating mismatches in the cloning primers. **USP13** (NM_003940) was synthesised by GenScript to incorporate an N terminal HA tag, two sites of siRNA resistance and loss of BamHI and BglII sites by synonymous mutations. The cDNA was cloned into BamHI-XhoI sites. The following plasmids were from Addgene FLAG-SENP1 (#17357, Edward Yeh (Cheng et al., 2007)) GFP-SENP3, GFP-SENP5 (#34554, #34555 Mary Dasso, (Yun et al., 2008)) and FLAG-SENP6 (#18065, Edward Yeh, (Dou et al., 2010)).

### Cell lines

The growth conditions and vendors for all cell lines are details in table 2. FlpIn stable cell lines were generated using HEK293^TrEx-FlpIn^ (Invitrogen) and HeLa FlpIn (a gift from Grant Stewart, University of Birmingham) cells transfected with pcDNA5/FRT/TO based vectors and the recombinase pOG44 (Invitrogen) using FuGene6 (Promega). After 48 hr, cells were placed into hygromycin selection media (100 μg/ml) and grown until colonies formed on plasmid-transfected plates but not controls. HAP1 SENP2 knockout cells (128bp deletion in exon 3, HZGHC002974c003) and parental cells were from Horizon Discovery and were cultured according to manufacturer’s instructions.

### Co-IP

HEK293^FlpIn^ myc-MDC1^WT^ were seeded on 10cm plates in the presence of doxycycline (1μg/mL) for 24 hr prior to transfection with FLAG-SENP2 (3μg / plate) for a further 48 hr. Cells were treated with 4 Gy IR and pelleted 1 hr later in cold PBS. Cell pellets were lysed in 0.5mL hypotonic buffer (10mM HEPES pH 7.8, 10mM KCl, 1.5mM MgCl_2_, 340mM Sucrose, 10% glycerol 0.2% NP40, protease and phosphatase inhibitor cocktails) for 5 minutes on ice and centrifuged at 3,000 rpm for 3 minutes. The nuclear pellet was lysed in nuclear buffer (0.05% NP40, 50mM Tris pH 8, 300mM NaCl, protease and phosphatase inhibitor cocktails) and rotated for 30 minutes at 4°C. Lysates were briefly sonicated and clarified at 12,000 rpm for 10 minutes to remove debris. Cleared lysates (0.9mL) were incubated with either myc (Thermo-Fisher) or M2 (Sigma) agarose (20μL packed bead volume) at 4°C with rotation for 16 hr. Beads were washed 3x with NETN buffer (100mM NaCl, 20mM Tris-HCl pH 8, 0.5mM EDTA and 0.5% NP40) before elution with 4X Lamelli buffer.

### His-SUMO Pulldown

HEK293^FlpIn^ 6xHis-myc-SUMO1 or SUMO2 were seeded on 10cm plates in the presence of doxycycline (1μg/mL) for 24 hr prior to knockdown with indicated siRNA for a further 48 hr. Cells were treated with 10 Gy IR and pelleted 1 hr later in cold PBS. Cell pellets were lysed in 8M Urea buffer (8⁗M urea, 0.1⁗M Na_2_HPO_4_/NaH_2_PO_4_, 0.01⁗M Tris–HCl, pH 6.3, 10⁗mM β-mercaptoethanol, 5⁗mM imidazole plus 0.2% Triton-X-100) with vigorous pipetting. Lysates were left on ice for 30 minutes prior to sonication and clarification at 12,000 rpm for 10 minutes. Cleared lysates (0.9mL) were incubated with Nickel-agarose (HIS-Select, Sigma) (30μL packed bead volume) at 4°C with rotation for 16 hr. Beads were washed 3x with 8M Urea buffer before elution with 4X Lamelli buffer.

### Metaphases

HeLa^FlpIn^ or HeLa^FlpIn^ SENP2^WT^ cells were plated on 60mm plates in the presence of doxycycline for 48 hr prior to irradiation at 2 Gy. Eighteen hr later cells were incubated with Colcemid (0.05 μg/ml) 6 hr. Cells were then trypsinized and centrifuged at 1200 rpm for 5 minutes. Supernatant was discarded and cells re-suspended. 5 ml of ice-cold 0.56% KCl solution was then added and incubated at room temperature for 15 min before centrifuging at 1200 rpm for 5 min. Supernatant was discarded and cell pellet broken before fixation. Cells were then fixed in 5 ml of ice-cold methanol: glacial acetic acid (3:1). Fixation agents were removed and 10 μl of cells suspension was dropped onto alcohol cleaned slide. Slides were allowed to dry at least 24 hr and then stained with Giemsa solution (Sigma) diluted 1:20 for 20 min. Slide mounting was performed with Eukitt (Sigma).

Overexpression and purification of MDC1^WT^ and MDC1^K1840R^ (aa 1818-2094) C-terminal domains.

The expression of His-SUMO MDC1^WT^ and His-SUMO-MDC1^K1840R^ in BL21(DE3*)/pCA528-MDC1 was induced by the addition of 1 mM Isopropyl-β-d-thiogalactopyranoside (IPTG), and the proteins were produced in LB medium containing 100 μg/ml of kanamycin overnight at 18°C. For purification of the His-SUMO MDC1^WT^ and His-SUMO MDC1^K1840R^ products, the cells were harvested and re-suspended in 20 mM HEPES potassium salt, pH 7.4, 50 mM Imidazole, 500 mM NaCl, 1.0 mM TCEP [tris(2-carboxyethyl)phosphine], complete EDTA-free protease inhibitor cocktail tablet (Roche). Cells were lysed using an Emulsiflex-C3 homogenizer (Avestin) and broken by three passages through the chilled cell. The lysate was centrifuged at 75,000 xg using a JA 25 rotor (Beckman Coulter) and filtered through a 0.45-μm filter. The clarified lysate was applied onto a 5-ml HisTrap HP column (GE Healthcare). The column was washed extensively using the same buffer, and the protein was eluted using buffer containing 500 mM imidazole.

Fractions containing a band of the correct size were concentrated using a Vivaspin 20-ml concentrator (10,000 molecular weight cut-off [MWCO]) (GE Healthcare) and gel purified using an Akta Pure 25 (GE Healthcare LS) with a prepacked Hi-Load 10/300 Superdex 200 PG column.

For removal of the His-SUMO tag, 1ul of ULP-1 (20mg/ml) was added to 5ml of His-SUMO MDC1^WT^ and His-SUMO-MDC1^K1840R^ and left overnight at 4°C. The samples were concentrated to 500μl using a Vivaspin 4-ml concentrator (10,000 molecular weight cut-off [MWCO]) (GE Healthcare) and gel purified on a Hi-Load 10/300 Superdex 75 PG column in order to separate the untagged proteins from the ULP-1 protease and the cleaved His-SUMO tag.

### *In vitro* SUMOylation assay

*In vitro* SUMOylation assay reactions were typically performed in a total volume of 20 μl with 200 ng recombinant Human SUMO E1 (SAE1/UBA2) (R&D Systems), 100 ng of Ubc9 (Boston Biochem), 1 µg of SUMO2, (Boston Biochem), 1 µg of recombinant untagged-MDC1 (aa1818–2094) and untagged MDC1^K1840R^. Reaction buffer (50 mM HEPES, 50 mM MgCl_2_, 0.5 mM DTT) was added to a final 1x concentration and supplemented with 4 mM ATP-Mg. Reactions were incubated at 30C for 1h and stopped by addition of 2x Laemmli loading buffer.

### *In vitro* deSUMOylation assayx

For de-SUMOylation; the *in vitro* SUMOylation reaction was split in two and SENP2 catalytic domain (Boston Biochem) was added to a final concentration of 50 nM. Reactions were incubated at 30°C for 0.5 hr and stopped by addition of 2x Laemmli loading buffer (Sigma).

### Statistics

Unless otherwise stated all statistical analysis was by two-sided Students T-test throughout. *<p0.05, **p<0.01, ***P<0.005 ****P<0.001. All centre values are given as the mean and all error bars are standard error about the mean (s.e.m). Data was analysed using GraphPad Prism 7.03.

### Quantification

All Western Blot or Image analysis for quantification was done using ImageJ unless otherwise specified.

## Supporting information

## Acknowledgements

Grant funding for this project was as follows. Wellcome Trust: AJG, AKW, ASC, MJ (206343/Z/17/Z), Cancer Research UK: C8820/A19062 AJG, HRS, RMD: Breast Cancer Now (2015MayPR499) KS, and CRUK Centre training MDM. In addition, we thank James Beesley for technical assistance, Jeremy Stark for the U2OS reporter cell lines (City of Hope, USA) and the Microscopy, Imaging and FACS services at Birmingham University in the Tech Hub facility for equipment support and maintenance.

## Author contributions

Cloning and generation of stable cell lines (AJG, AW and RMD), Co-IP and pulldowns (AJG). IF analysis (AJG, AW, MDM, HRS), FACS analysis (AW and RMD), Metaphase spreads (KS). Immunoblots (AJG, AW, ASC, HM). The paper was written and project conceived by AJG and JRM. All authors reviewed the manuscript.

## References

Bologna, S., Altmannova, V., Valtorta, E., Koenig, C., Liberali, P., Gentili, C., Anrather, D., Ammerer, G., Pelkmans, L., Krejci, L., et al. (2015). Sumoylation regulates EXO1 stability and processing of DNA damage. Cell Cycle 14, 2439–2450.

Butler, L.R., Densham, R.M., Jia, J., Garvin, A.J., Stone, H.R., Shah, V., Weekes, D., Festy, F., Beesley, J., and Morris, J.R. (2012). The proteasomal de-ubiquitinating enzyme POH1 promotes the double-strand DNA break response. The EMBO journal 31, 3918–3934.

Cancer Genome Atlas, N. (2015). Comprehensive genomic characterization of head and neck squamous cell carcinomas. Nature 517, 576–582.

Castillo-Villanueva, E., Ballesteros, G., Schmid, M., Hidalgo, P., Schreiner, S., Dobner, T., and Gonzalez, R.A. (2014). The Mre11 Cellular Protein Is Modified by Conjugation of Both SUMO-1 and SUMO-2/3 during Adenovirus Infectio. ISRN Virology 2014.

Chapman, J.R., and Jackson, S.P. (2008). Phospho-dependent interactions between NBS1 and MDC1 mediate chromatin retention of the MRN complex at sites of DNA damage. EMBO Rep 9, 795–801.

Chen, Y.J., Chuang, Y.C., Chuang, C.N., Cheng, Y.H., Chang, C.R., Leng, C.H., and Wang, T.F. (2016). S. cerevisiae Mre11 recruits conjugated SUMO moieties to facilitate the assembly and function of the Mre11-Rad50-Xrs2 complex. Nucleic Acids Res 44, 2199–2213.

Cheng, J., Kang, X.L., Zhang, S., and Yeh, E.T.H. (2007). SUMO-Specific protease 1 is essential for stabilization of HIF1 alpha during hypoxia. Cell 131, 584–595.

Chow, K.H., Elgort, S., Dasso, M., Powers, M.A., and Ullman, K.S. (2014). The SUMO proteases SENP1 and SENP2 play a critical role in nucleoporin homeostasis and nuclear pore complex function. Mol Biol Cell 25, 160–168.

Chroma, K., Mistrik, M., Moudry, P., Gursky, J., Liptay, M., Strauss, R., Skrott, Z., Vrtel, R., Bartkova, J., Kramara, J., et al. (2017). Tumors overexpressing RNF168 show altered DNA repair and responses to genotoxic treatments, genomic instability and resistance to proteotoxic stress. Oncogene 36, 2405–2422.

Danielsen, J.R., Povlsen, L.K., Villumsen, B.H., Streicher, W., Nilsson, J., Wikstrom, M., Bekker-Jensen, S., and Mailand, N. (2012). DNA damage-inducible SUMOylation of HERC2 promotes RNF8 binding via a novel SUMO-binding Zinc finger. J Cell Biol 197, 179–187.

Dantuma, N.P., Acs, K., and Luijsterburg, M.S. (2014). Should I stay or should I go: VCP/p97-mediated chromatin extraction in the DNA damage response. Experimental Cell Research 329, 9–17.

Doil, C., Mailand, N., Bekker-Jensen, S., Menard, P., Larsen, D.H., Pepperkok, R., Ellenberg, J., Panier, S., Durocher, D., Bartek, J., et al. (2009). RNF168 Binds and Amplifies Ubiquitin Conjugates on Damaged Chromosomes to Allow Accumulation of Repair Proteins. Cell 136, 435–446.

Dou, H., Huang, C., Singh, M., Carpenter, P.B., and Yeh, E.T.H. (2010). Regulation of DNA Repair through DeSUMOylation and SUMOylation of Replication Protein A Complex. Molecular Cell 39, 333– 345.

Eladad, S., Ye, T.Z., Hu, P., Leversha, M., Beresten, S., Matunis, M.J., and Ellis, N.A. (2005). Intra-nuclear trafficking of the BLM helicase to DNA damage-induced foci is regulated by SUMO modification. Hum Mol Genet 14, 1351–1365.

Galanty, Y., Belotserkovskaya, R., Coates, J., and Jackson, S.P. (2012). RNF4, a SUMO-targeted ubiquitin E3 ligase, promotes DNA double-strand break repair. Genes Dev 26, 1179–1195.

Galanty, Y., Belotserkovskaya, R., Coates, J., Polo, S., Miller, K.M., and Jackson, S.P. (2009). Mammalian SUMO E3-ligases PIAS1 and PIAS4 promote responses to DNA double-strand breaks. Nature 462, 935–U132.

Garvin, A.J., Densham, R., Blair-Reid, S.A., Pratt, K.M., Stone, H.R., Weekes, D., Lawrence, K.J., and Morris, J.R. (2013). The deSUMOylase SENP7 promotes chromatin relaxation for homologous recombination DNA repair. EMBO Reports 14, 975–983.

Garvin, A.J., and Morris, J.R. (2017). SUMO, a small, but powerful, regulator of double-strand break repair. Philos Trans R Soc Lond B Biol Sci 372.

Goeres, J., Chan, P.K., Mukhopadhyay, D., Zhang, H., Raught, B., and Matunis, M.J. (2011). The SUMO-specific isopeptidase SENP2 associates dynamically with nuclear pore complexes through interactions with karyopherins and the Nup107-160 nucleoporin subcomplex. Mol Biol Cell 22, 4868– 4882.

Hang, J., and Dasso, M. (2002). Association of the human SUMO-1 protease SENP2 with the nuclear pore. J Biol Chem 277, 19961–19966.

Hang, L.E., Lopez, C.R., Liu, X., Williams, J.M., Chung, I., Wei, L., Bertuch, A.A., and Zhao, X. (2014). Regulation of Ku-DNA association by Yku70 C-terminal tail and SUMO modification. J Biol Chem 289, 10308–10317.

He, X., Riceberg, J., Soucy, T., Koenig, E., Minissale, J., Gallery, M., Bernard, H., Yang, X., Liao, H., Rabino, C., et al. (2017). Probing the roles of SUMOylation in cancer cell biology by using a selective SAE inhibitor. Nat Chem Biol 13, 1164–1171.

Hecker, C.M., Rabiller, M., Haglund, K., Bayer, P., and Dikic, I. (2006). Specification of SUMO1- and SUMO2-interacting motifs. J Biol Chem 281, 16117–16127.

Hendriks, I.A., Lyon, D., Su, D., Skotte, N.H., Daniel, J.A., Jensen, L.J., and Nielsen, M.L. (2018). Site-specific characterization of endogenous SUMOylation across species and organs. Nat Commun 9, 2456.

Hendriks, I.A., Treffers, L.W., Verlaan-de Vries, M., Olsen, J.V., and Vertegaal, A.C. (2015). SUMO-2 Orchestrates Chromatin Modifiers in Response to DNA Damage. Cell Rep.

Hendriks, I.A., and Vertegaal, A.C. (2015). SUMO in the DNA damage response. Oncotarget 6, 15734– 15735.

Hendriks, I.A., and Vertegaal, A.C. (2016). A comprehensive compilation of SUMO proteomics. Nat Rev Mol Cell Biol 17, 581–595.

Ismail, I.H., Gagne, J.P., Caron, M.C., McDonald, D., Xu, Z., Masson, J.Y., Poirier, G.G., and Hendzel, M.J. (2012). CBX4-mediated SUMO modification regulates BMI1 recruitment at sites of DNA damage. Nucleic acids research 40, 5497–5510.

Jentsch, S., and Psakhye, I. (2013). Control of nuclear activities by substrate-selective and protein-group SUMOylation. Annu Rev Genet 47, 167–186.

Kessler, J.D., Kahle, K.T., Sun, T., Meerbrey, K.L., Schlabach, M.R., Schmitt, E.M., Skinner, S.O., Xu, Q., Li, M.Z., Hartman, Z.C., et al. (2012). A SUMOylation-dependent transcriptional subprogram is required for Myc-driven tumorigenesis. Science 335, 348–353.

Kumar, A., Ito, A., Hirohama, M., Yoshida, M., and Zhang, K.Y. (2016). Identification of new SUMO activating enzyme 1 inhibitors using virtual screening and scaffold hopping. Bioorg Med Chem Lett 26, 1218–1223.

Kumar, A., Ito, A., Takemoto, M., Yoshida, M., and Zhang, K.Y. (2014). Identification of 1,2,5-oxadiazoles as a new class of SENP2 inhibitors using structure based virtual screening. J Chem Inf Model 54, 870–880.

Kumar, R., and Cheok, C.F. (2017). Dynamics of RIF1 SUMOylation is regulated by PIAS4 in the maintenance of Genomic Stability. Sci Rep 7, 17367.

Kumar, R., Gonzalez-Prieto, R., Xiao, Z., Verlaan-de Vries, M., and Vertegaal, A.C.O. (2017). The STUbL RNF4 regulates protein group SUMOylation by targeting the SUMO conjugation machinery. Nat Commun 8, 1809.

Kung, C.C., Naik, M.T., Wang, S.H., Shih, H.M., Chang, C.C., Lin, L.Y., Chen, C.L., Ma, C., Chang, C.F., and Huang, T.H. (2014). Structural analysis of poly-SUMO chain recognition by the RNF4-SIMs domain. Biochem J 462, 53–65.

Lamoliatte, F., Caron, D., Durette, C., Mahrouche, L., Maroui, M.A., Caron-Lizotte, O., Bonneil, E., Chelbi-Alix, M.K., and Thibault, P. (2014). Large-scale analysis of lysine SUMOylation by SUMO remnant immunoaffinity profiling. Nat Commun 5, 5409.

Lee, M.H., Mabb, A.M., Gill, G.B., Yeh, E.T., and Miyamoto, S. (2011). NF-kappaB induction of the SUMO protease SENP2: A negative feedback loop to attenuate cell survival response to genotoxic stress. Molecular Cell 43, 180–191.

Li, Y., Luo, K., Yin, Y., Wu, C., Deng, M., Li, L., Chen, Y., Nowsheen, S., Lou, Z., and Yuan, J. (2017). USP13 regulates the RAP80-BRCA1 complex dependent DNA damage response. Nat Commun 8, 15752.

Li, Y.J., Stark, J.M., Chen, D.J., Ann, D.K., and Chen, Y. (2010). Role of SUMO:SIM-mediated protein-protein interaction in non-homologous end joining. Oncogene 29, 3509–3518.

Licciardello, M.P., Mullner, M.K., Durnberger, G., Kerzendorfer, C., Boidol, B., Trefzer, C., Sdelci, S., Berg, T., Penz, T., Schuster, M., et al. (2015). NOTCH1 activation in breast cancer confers sensitivity to inhibition of SUMOylation. Oncogene 34, 3780–3790.

Lok, G.T., Sy, S.M., Dong, S.S., Ching, Y.P., Tsao, S.W., Thomson, T.M., and Huen, M.S. (2011). Differential regulation of RNF8-mediated Lys48- and Lys63-based poly-ubiquitylation. Nucleic acids research.

Lu, X., Olsen, S.K., Capili, A.D., Cisar, J.S., Lima, C.D., and Tan, D.S. (2010). Designed semisynthetic protein inhibitors of Ub/Ubl E1 activating enzymes. J Am Chem Soc 132, 1748–1749.

Luo, K., Zhang, H., Wang, L., Yuan, J., and Lou, Z. (2012). Sumoylation of MDC1 is important for proper DNA damage response. The EMBO journal 31, 3008–3019.

Madu, I.G., Namanja, A.T., Su, Y., Wong, S., Li, Y.J., and Chen, Y. (2013). Identification and characterization of a new chemotype of noncovalent SENP inhibitors. ACS Chem Biol 8, 1435–1441.

Makhnevych, T., Ptak, C., Lusk, C.P., Aitchison, J.D., and Wozniak, R.W. (2007). The role of karyopherins in the regulated sumoylation of septins. J Cell Biol 177, 39–49.

Morris, J.R., Boutell, C., Keppler, M., Densham, R., Weekes, D., Alamshah, A., Butler, L., Galanty, Y., Pangon, L., Kiuchi, T., et al. (2009). The SUMO modification pathway is involved in the BRCA1 response to genotoxic stress. Nature 462, 886–U877.

Morris, J.R., and Garvin, A.J. (2017). SUMO in the DNA Double-Stranded Break Response: Similarities, Differences, and Cooperation with Ubiquitin. J Mol Biol 429, 3376–3387.

Mukhopadhyay, D., and Dasso, M. (2007). Modification in reverse: the SUMO proteases. Trends Biochem Sci 32, 286–295.

Nishi, R., Wijnhoven, P., le Sage, C., Tjeertes, J., Galanty, Y., Forment, J.V., Clague, M.J., Urbe, S., and Jackson, S.P. (2014). Systematic characterization of deubiquitylating enzymes for roles in maintaining genome integrity. Nat Cell Biol 16, 1016-1026, 1011-1018.

Nowsheen, S., Aziz, K., Aziz, A., Deng, M., Qin, B., Luo, K., Jeganathan, K.B., Zhang, H., Liu, T., Yu, J., et al. (2018). L3MBTL2 orchestrates ubiquitin signalling by dictating the sequential recruitment of RNF8 and RNF168 after DNA damage. Nat Cell Biol 20, 455–464.

Odeh, H.M., Coyaud, E., Raught, B., and Matunis, M.J. (2018). The SUMO-Specific Isopeptidase SENP2 is Targeted to Intracellular Membranes via a Predicted N-Terminal Amphipathic alpha-Helix. Mol Biol Cell, mbcE17070445.

Ouyang, K.J., Woo, L.L., Zhu, J., Huo, D., Matunis, M.J., and Ellis, N.A. (2009). SUMO modification regulates BLM and RAD51 interaction at damaged replication forks. PLoS Biol 7, e1000252.

Panier, S., and Boulton, S.J. (2014). Double-strand break repair: 53BP1 comes into focus. Nat Rev Mol Cell Biol 15, 7–18.

Panse, V.G., Kuster, B., Gerstberger, T., and Hurt, E. (2003). Unconventional tethering of Ulp1 to the transport channel of the nuclear pore complex by karyopherins. Nat Cell Biol 5, 21–27.

Pfeiffer, A., Luijsterburg, M.S., Acs, K., Wiegant, W.W., Helfricht, A., Herzog, L.K., Minoia, M., Bottcher, C., Salomons, F.A., van Attikum, H., et al. (2017). Ataxin-3 consolidates the MDC1-dependent DNA double-strand break response by counteracting the SUMO-targeted ubiquitin ligase RNF4. EMBO J 36, 1066–1083.

Plechanovova, A., Jaffray, E.G., Tatham, M.H., Naismith, J.H., and Hay, R.T. (2012). Structure of a RING E3 ligase and ubiquitin-loaded E2 primed for catalysis. Nature 489, 115–U135.

Poulsen, S.L., Hansen, R.K., Wagner, S.A., van Cuijk, L., van Belle, G.J., Streicher, W., Wikstrom, M., Choudhary, C., Houtsmuller, A.B., Marteijn, J.A., et al. (2013). RNF111/Arkadia is a SUMO-targeted ubiquitin ligase that facilitates the DNA damage response. J Cell Biol 201, 797–807.

Psakhye, I., and Jentsch, S. (2012). Protein Group Modification and Synergy in the SUMO Pathway as Exemplified in DNA Repair. Cell 151, 807–820.

Qian, J., and Massion, P.P. (2008). Role of chromosome 3q amplification in lung cancer. J Thorac Oncol 3, 212–215.

Rojas-Fernandez, A., Plechanovova, A., Hattersley, N., Jaffray, E., Tatham, M.H., and Hay, R.T. (2014). SUMO chain-induced dimerization activates RNF4. Mol Cell 53, 880–892.

Savic, V., Yin, B., Maas, N.L., Bredemeyer, A.L., Carpenter, A.C., Helmink, B.A., Yang-Iott, K.S., Sleckman, B.P., and Bassing, C.H. (2009). Formation of dynamic gamma-H2AX domains along broken DNA strands is distinctly regulated by ATM and MDC1 and dependent upon H2AX densities in chromatin. Mol Cell 34, 298–310.

Seeler, J.S., and Dejean, A. (2017). SUMO and the robustness of cancer. Nat Rev Cancer 17, 184–197.

Shao, G., Lilli, D.R., Patterson-Fortin, J., Coleman, K.A., Morrissey, D.E., and Greenberg, R.A. (2009). The Rap80-BRCC36 de-ubiquitinating enzyme complex antagonizes RNF8-Ubc13-dependent ubiquitination events at DNA double strand breaks. Proceedings of the National Academy of Sciences of the United States of America 106, 3166–3171.

Shima, H., Suzuki, H., Sun, J.Y., Kono, K., Shi, L., Kinomura, A., Horikoshi, Y., Ikura, T., Ikura, M., Kanaar, R., et al. (2013). Activation of the SUMO modification system is required for the accumulation of RAD51 at sites of DNA damage. Journal of Cell Science 126, 5284–5292.

Song, J., Durrin, L.K., Wilkinson, T.A., Krontiris, T.G., and Chen, Y. (2004). Identification of a SUMO-binding motif that recognizes SUMO-modified proteins. Proc Natl Acad Sci U S A 101, 14373–14378.

Stewart, G.S., Panier, S., Townsend, K., Al-Hakim, A.K., Kolas, N.K., Miller, E.S., Nakada, S., Ylanko, J., Olivarius, S., Mendez, M., et al. (2009). The RIDDLE syndrome protein mediates a ubiquitin-dependent signaling cascade at sites of DNA damage. Cell 136, 420–434.

Stucki, M., Clapperton, J.A., Mohammad, D., Yaffe, M.B., Smerdon, S.J., and Jackson, S.P. (2005). MDC1 directly binds phosphorylated histone H2AX to regulate cellular responses to DNA double-strand breaks. Cell 123, 1213–1226.

Tammsalu, T., Matic, I., Jaffray, E.G., Ibrahim, A.F., Tatham, M.H., and Hay, R.T. (2014). Proteome-wide identification of SUMO2 modification sites. Science signaling 7, rs2.

Tan, M., Gong, H., Wang, J., Tao, L., Xu, D., Bao, E., Liu, Z., and Qiu, J. (2015). SENP2 regulates MMP13 expression in a bladder cancer cell line through SUMOylation of TBL1/TBLR1. Sci Rep 5, 13996.

Tatham, M.H., Geoffroy, M.C., Shen, L., Plechanovova, A., Hattersley, N., Jaffray, E.G., Palvimo, J.J., and Hay, R.T. (2008). RNF4 is a poly-SUMO-specific E3 ubiquitin ligase required for arsenic-induced PML degradation. Nat Cell Biol 10, 538–546.

Thorslund, T., Ripplinger, A., Hoffmann, S., Wild, T., Uckelmann, M., Villumsen, B., Narita, T., Sixma, T.K., Choudhary, C., Bekker-Jensen, S., et al. (2015). Histone H1 couples initiation and amplification of ubiquitin signalling after DNA damage. Nature 527, 389-+.

Torrecilla, I., Oehler, J., and Ramadan, K. (2017). The role of ubiquitin-dependent segregase p97 (VCP or Cdc48) in chromatin dynamics after DNA double strand breaks. Philos Trans R Soc Lond B Biol Sci 372.

Uckelmann, M., and Sixma, T.K. (2017). Histone ubiquitination in the DNA damage response. DNA Repair (Amst).

Vyas, R., Kumar, R., Clermont, F., Helfricht, A., Kalev, P., Sotiropoulou, P., Hendriks, I.A., Radaelli, E., Hochepied, T., Blanpain, C., et al. (2013). RNF4 is required for DNA double-strand break repair in vivo. Cell Death Differ 20, 490–502.

Yin, Y.L., Seifert, A., Chua, J.S., Maure, J.F., Golebiowski, F., and Hay, R.T. (2012). SUMO-targeted ubiquitin E3 ligase RNF4 is required for the response of human cells to DNA damage. Genes & Development 26, 1196–1208.

Yu, B., Swatkoski, S., Holly, A., Lee, L.C., Giroux, V., Lee, C.S., Hsu, D., Smith, J.L., Yuen, G., Yue, J., et al. (2015). Oncogenesis driven by the Ras/Raf pathway requires the SUMO E2 ligase Ubc9. Proc Natl Acad Sci U S A 112, E1724–1733.

Yun, C., Wang, Y., Mukhopadhyay, D., Backlund, P., Kolli, N., Yergey, A., Wilkinson, K.D., and Dasso, M. (2008). Nucleolar protein B23/nucleophosmin regulates the vertebrate SUMO pathway through SENP3 and SENP5 proteases. J Cell Biol 183, 589–595.

Yurchenko, V., Xue, Z., Gama, V., Matsuyama, S., and Sadofsky, M.J. (2008). Ku70 is stabilized by increased cellular SUMO. Biochem Biophys Res Commun 366, 263–268.

Yurchenko, V., Xue, Z., and Sadofsky, M.J. (2006). SUMO modification of human XRCC4 regulates its localization and function in DNA double-strand break repair. Mol Cell Biol 26, 1786–1794.

Zhang, H., Saitoh, H., and Matunis, M.J. (2002). Enzymes of the SUMO modification pathway localize to filaments of the nuclear pore complex. Mol Cell Biol 22, 6498–6508.

