## Supplementary material for "The deSUMOylase SENP2 coordinates homologous recombination and non-homologous end joining by independent mechanisms"

**Supplemental Data**

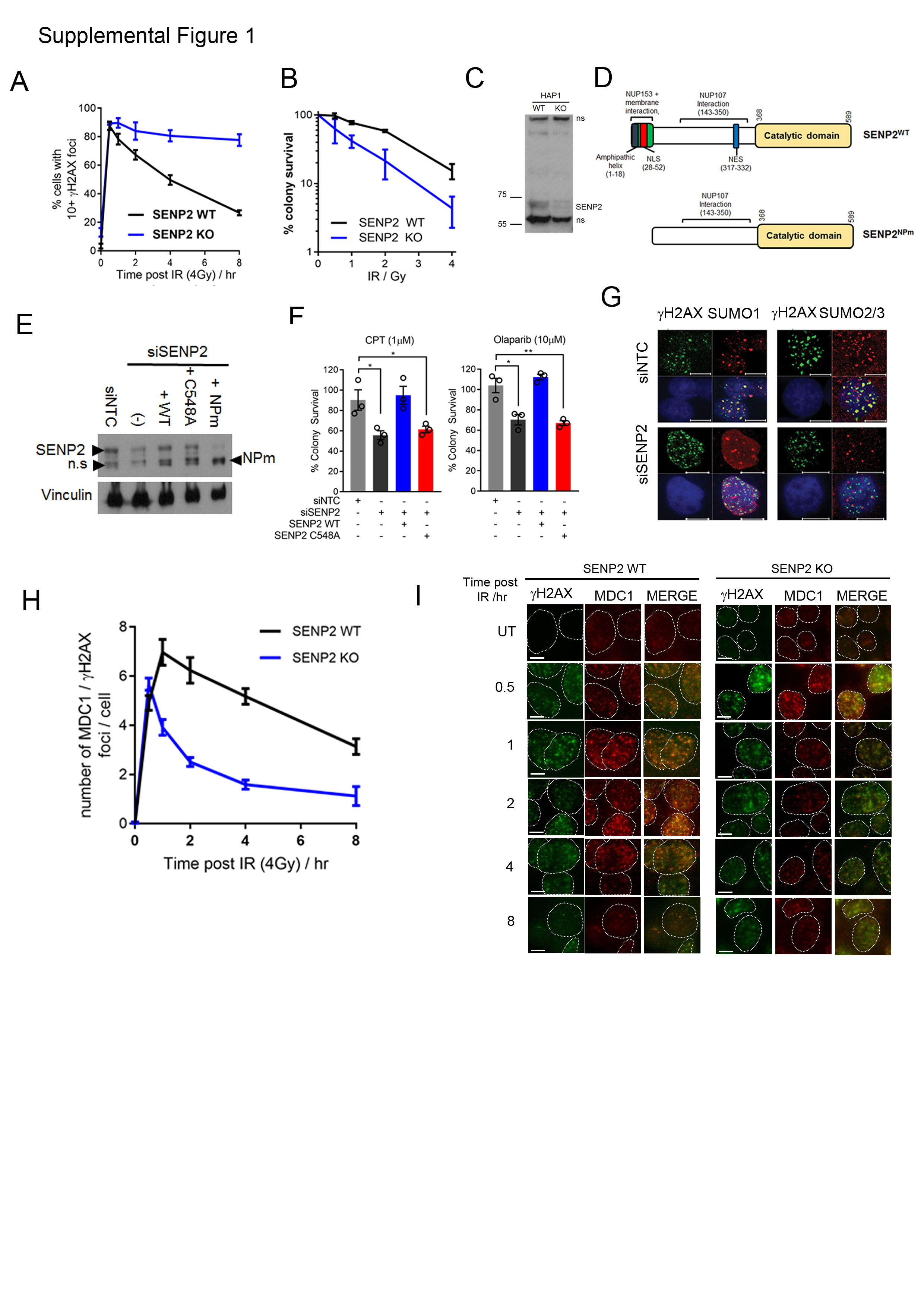

**Supplemental Figure 1. Related to Figure 1.**

**SENP2 promotes DNA damage signalling and DNA repair.**

**A.** Quantification of γH2AX foci in WT or *SENP2* KO HAP1 post 4 Gy IR. n=100 cells from 3 experiments.

**B**. IR colony survival of *SENP2-*KO and parental HAP1 cells, n=3

**C.** Western blot related to figure 1A. Note the SENP2^NPm^ migrates at a similar molecular weight to a lower band that cross reacts with the SENP2 antibody.­

**D.** Western blots of SENP2 protein expression levels in HAP1.

**E.** Cartoon schematic of human SENP2 domain locations, the N terminal 65 amino acids encompass the NLS (nuclear localisation signal) which also directs binding to the NUP153 nuclear pore component (Zhang, Saitoh, and Matunis 2002). An amphipathic helix between amino acids 1-18 directs SENP2 to cellular membranes (Odeh et al. 2018). The NES (nuclear export signal) directs SENP2 shuttling between the nucleus and cytoplasm. Interaction with NUP107 is through amino acids 143-350 although this interaction has not been as finely mapped as for NUP153. The NP mutant of SENP2 is illustrated below.

**F.** Colony survival as for 1A but using CPT (1 μM) or Olaparib (10 μM) both for 2 hr n=3.

**G.** IF images related to figure 1D. Scale bar = 10 μm

**H.** Quantification of MDC1 /γH2AX foci in HAP1 cells fixed at indicated times post 4 Gy IR, n= 100 cells.

**I.** Representative images related to S1H. Note HAP1 have smaller nuclei. Scale bar = 5 μm.

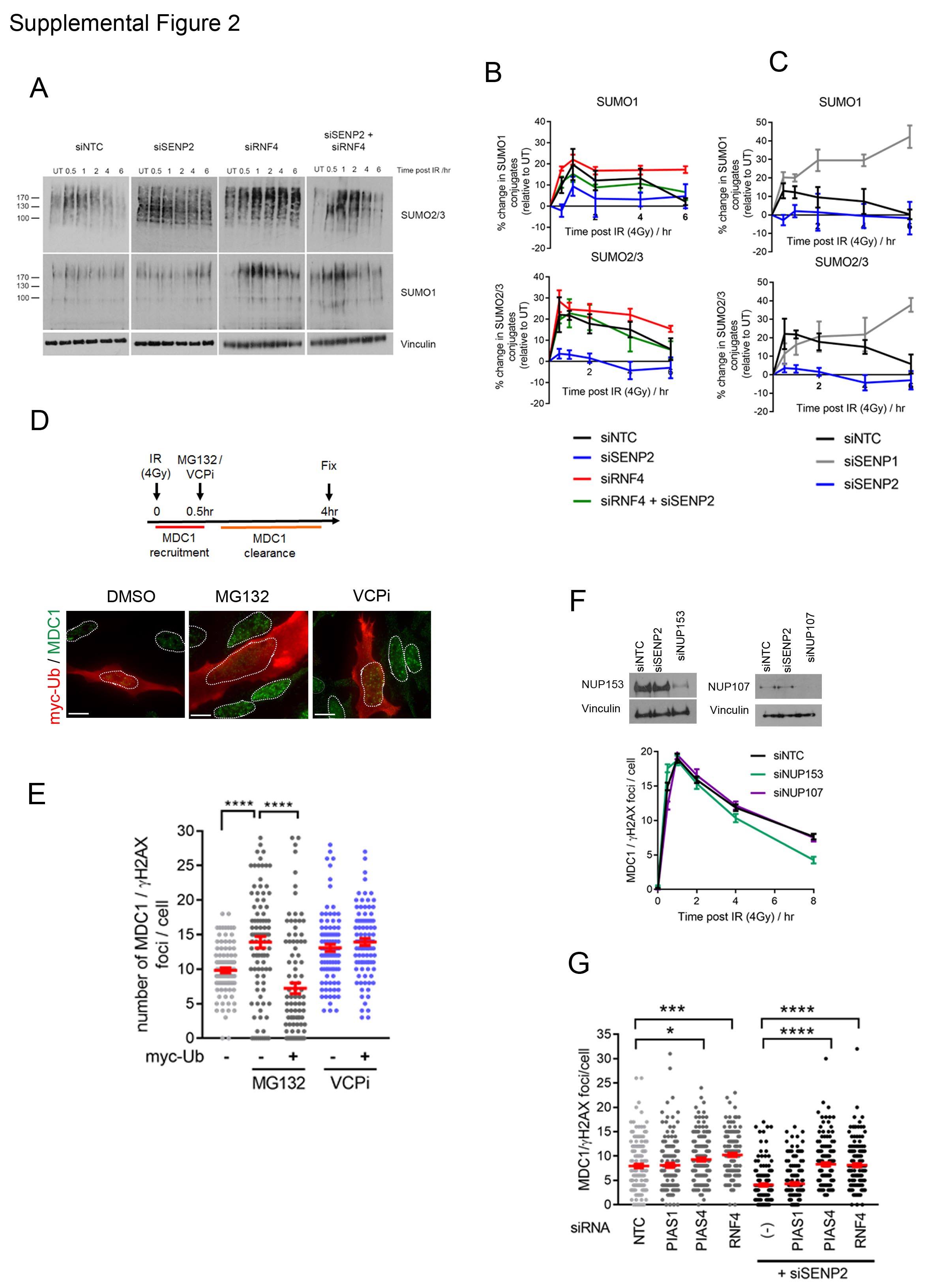

**Supplemental Figure 2. Related to Figure 1.**

**RNF4-VCP is responsible for defective DNA damage signalling in SENP2 depleted cells**

**A**. Western blots of SUMO1 and SUMO2/3 in HeLa^FlpIn^ treated with siRNA as indicated at various time points following 4 Gy IR.

**B-C.** Quantification of % change in SUMO conjugates (relative to non-irradiated SUMO conjugates) following 4 Gy IR related to S2A n=4

**D.** Top; cartoon of workflow of S2E, bottom; representative images related to S2E.

**E.** HeLa^FlpIn^ transfected with myc-ubiquitin and depleted with siNTC or siSENP2, irradiated with 4 Gy, 0.5 hr later cells were treated with DMSO or VCPi CB-5083 (0.1μM). Cells were fixed at 4 hours and scored for MDC1 foci / cell, n=100 from 3 experiments.

**F.** Quantification of MDC1 co-localising with γH2AX over time after 4 Gy IR in cells treated with indicated siRNAs. Top shows western blots to indicate knockdown, n=100

**G.** MDC1 / γH2AX foci quantification in HeLa depleted with indicated siRNA for 72 hr prior to irradiation (4 Gy) and fixation (4 hr post IR), n=100 cells.

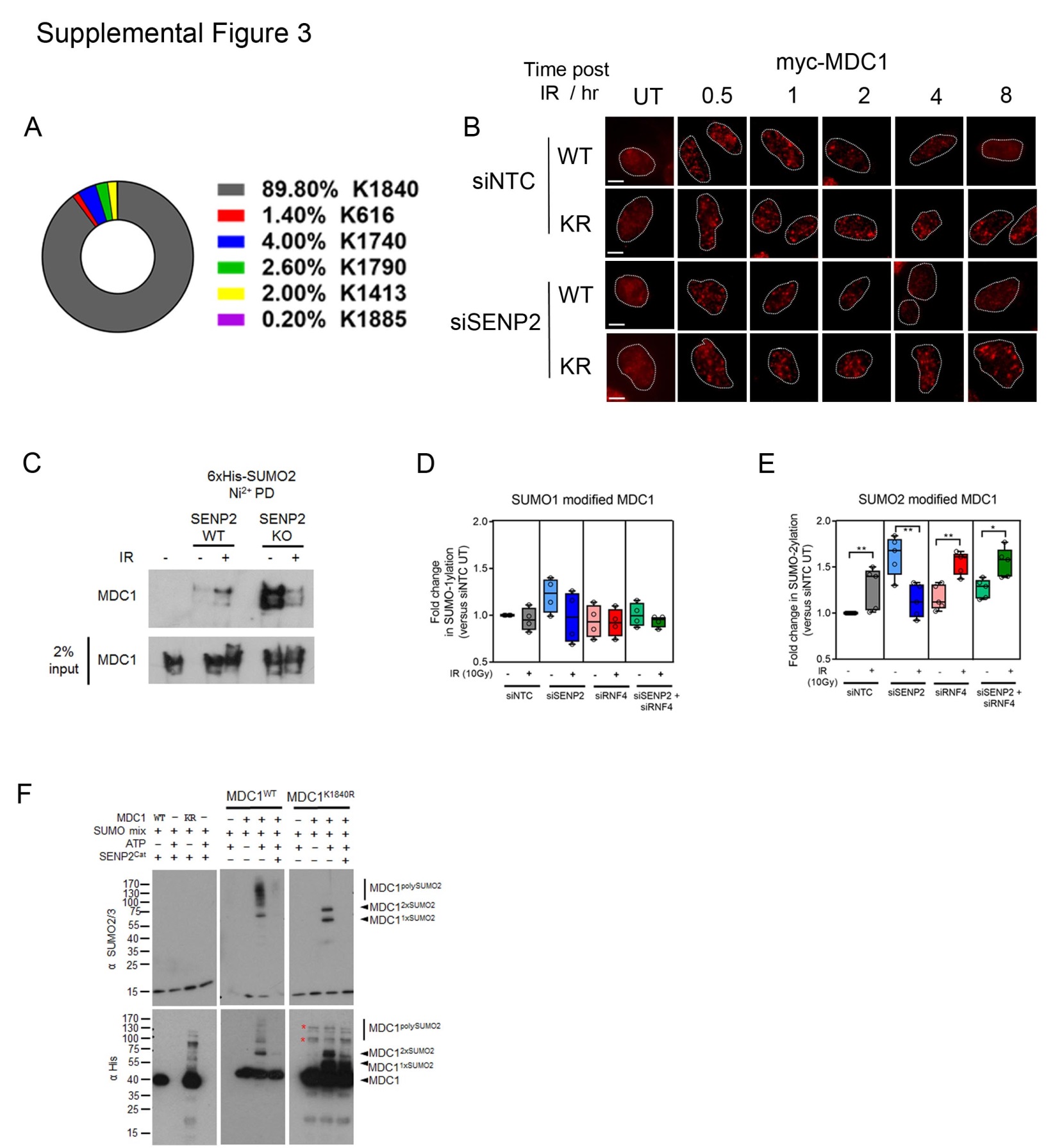

**Supplemental Figure 3. Related to Figure 2.**

**MDC1 is a SENP2 substrate and hypo-SUMOylation of MDC1 permits DDR signalling.**

**A**. Prevalence of detected SUMO conjugation sites on MDC1 given as fraction protein intensity as described in (Hendriks et al. 2017)

**B**. Related to figure 2A. HeLa^FlpIn^ myc-MDC1^WT^ or MDC1^K1840R^ cells transfected with either siNTC or siSENP2 and induced with the addition of Dox for 72 hours prior to irradiation (4 Gy). Cells were fixed at indicated times. Cells are immunostained for myc. Scale bar = 10 μm.

**C.** Ni^2+^ pulldown in HAP1 WT or SENP2 KO cells transiently transfected with 6xHis-myc-SUMO2 for 72 hours prior to irradiation (10 Gy). Western blots are probed with MDC1 antibody.

**D**. Quantification of the MDC1 purified by Ni^2+^ pulldowns in HEK293^FlpIn^ 6xHis-myc-SUMO1 cells treated with siRNAs as shown and either untreated or treated with 10 Gy IR and allowed to recover for 1 hour before harvesting. Relative enrichment in SUMO conjugates was determined by densitometry with the untreated siNTC sample being set as 1 (4 experiments). Error bars show s.e.m.

**E.** As for (S3D) but using HEK293^FlpIn^ 6xHis-myc-SUMO2 cells, n=5.

**F.** Recombinant His-MDC1^1818-2094^ fragments were SUMOylated *in vitro* with SUMO2, the product was divided in half with one half incubated with SENP2 catalytic domain for 30 minutes. Reactions were stopped with 2X Lamelli buffer, divided in half again and separated by SDS-PAGE. Reaction products were probed with His to detect MDC1 and SUMO2/3 to detect SUMOylated products. SUMO mix contains SUMO E1 and E2 enzymes. * denotes non-specific contaminants of the MDC1^K1840R^ fragment

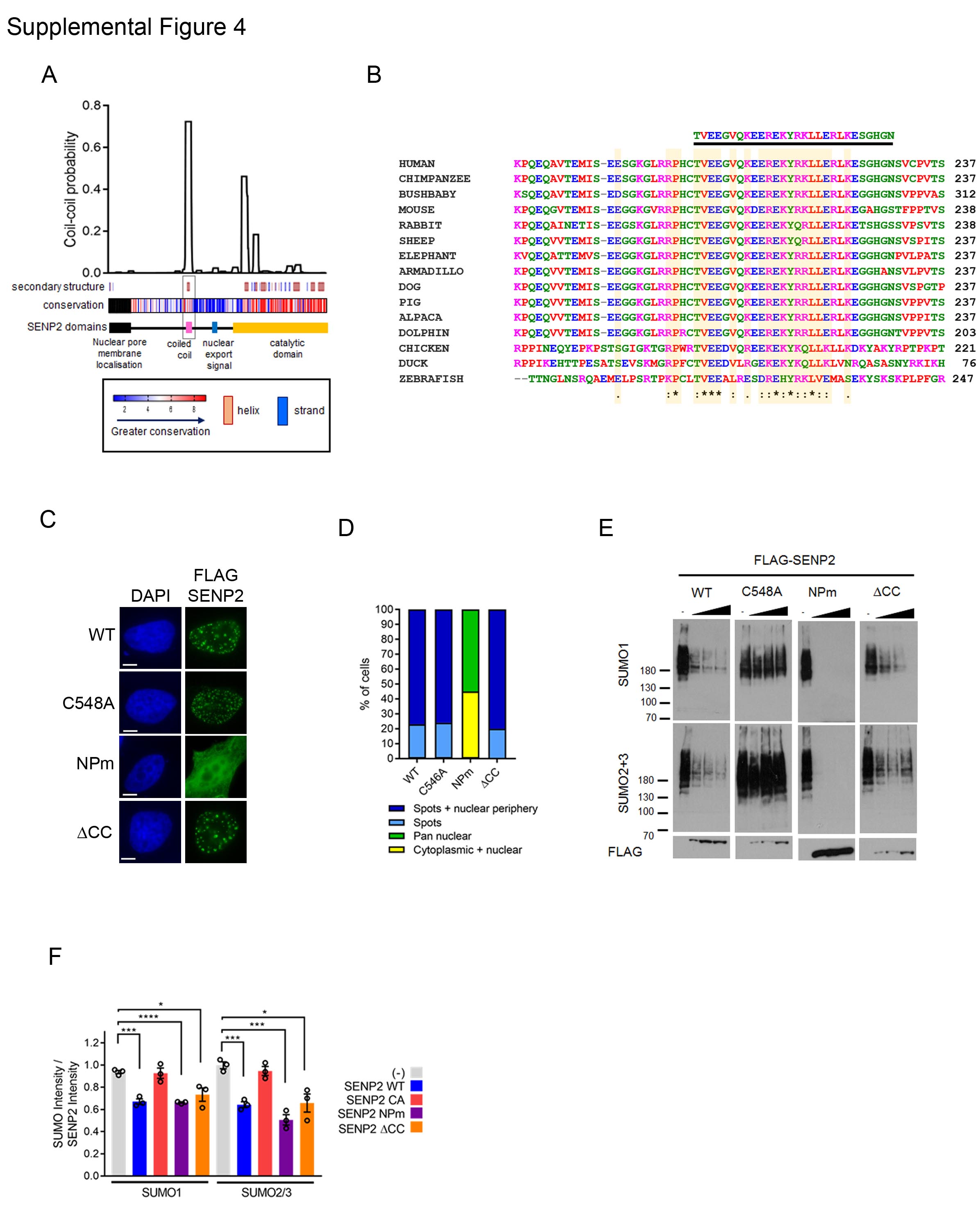

**Supplemental Figure 4. Related to Figure 2.**

**A conserved coiled-coil region of SENP2 contributes to MDC1 regulation**

**A.** Prediction of SENP2 coiled coil domain using NPS@ coiled coil prediction (Combet et al. 2000) using the MTIDK scoring matrix with no weight for the 2.5 patterns a and d. Data shown is the coil-coil probability with a 14 amino acid window. Predicted CCs that map to the catalytic domain were ignored as they identify as known helices. Secondary structure output was from the PredictProtein server (Rost, Yachdav, and Liu 2004). Conservation is shown as a heatmap using the 1-9 scores from Consurf (Ashkenazy et al. 2016) and higher amino acid conservation is shown in red. The black region in the N terminus was unscored as too few SENP2 species contain this region.

**B.** Sequence alignment of SENP2 using ClustalOmega centring on human SENP2 amino acids 178-237.

**C.** Representative localisation of FLAG - SENP2 variants in HeLa.

**D.** Quantification of SENP2 localisation in 100 cells.

**E.** SUMO1 and SUMO2/3 high molecular weight (HMW) conjugates in FLAG-SENP2 expressing HEK293 cells. FLAG-SENP2 was titrated to determine relative effects on SUMO conjugates versus SENP2 expression levels.

**F.** Quantification of (**E**). The intensity of the HMW SUMO conjugates was divided by the expression of FLAG-SENP2 to account for differences in expression levels. This was then shown relative to the intensity of SUMO HMW conjugates in mock transfected cells (set at 1.0). Data from 3 experiments.

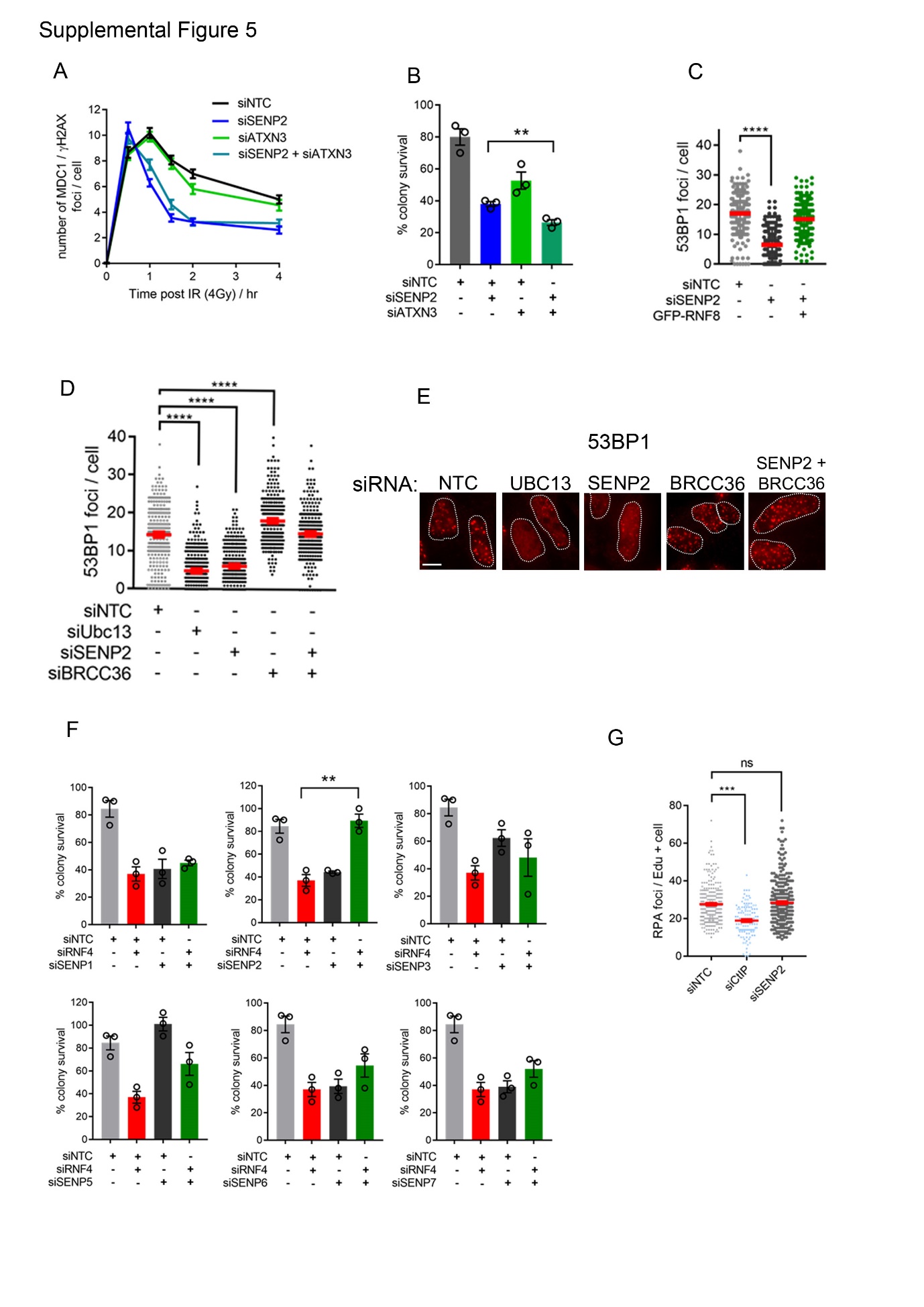

**Supplemental Figure 5. Related to Figure 3.**

**Requirement for SENP2 can be bypassed by increased K63-Ub signalling.**

**A.** Quantification of MDC1 foci co-localising with γH2AX staining in HeLa^FlpIn^ depleted with indicated siRNA. n=100 cells from 3 technical repeats. Graph shows mean number of co-localising foci per cell error bars are s.e.m.

**B.** IR colony survival of cells treated with indicated siRNA for 72 hours prior to irradiation (2 Gy). n=3

**C.** Quantification of 53BP1 foci 2 hours after 4 Gy IR in cells treated with the siRNAs shown and transfected with GFP-RNF8, n=50.

**D.** Quantification of 53BP1 foci 2 hours after 4 Gy IR in cells treated with the siRNAs shown n=100

**E.** Representative images related to (D). Scale bar = 10 μm

**F.** IR colony survival of a cells treated with indicated siRNA for 72 hours prior to irradiation (2 Gy), n=3.

**G.** RPA foci number in HeLa^FlpIn^ treated with indicated siRNA for 72 hours prior to irradiation (4 Gy) and 2 hour recovery. Cells were pulsed with 10 μM EdU 30 minutes prior to irradiation. The number of RPA70 foci per EdU positive cell is shown for a minimum of 100 cells across three independent experiments.

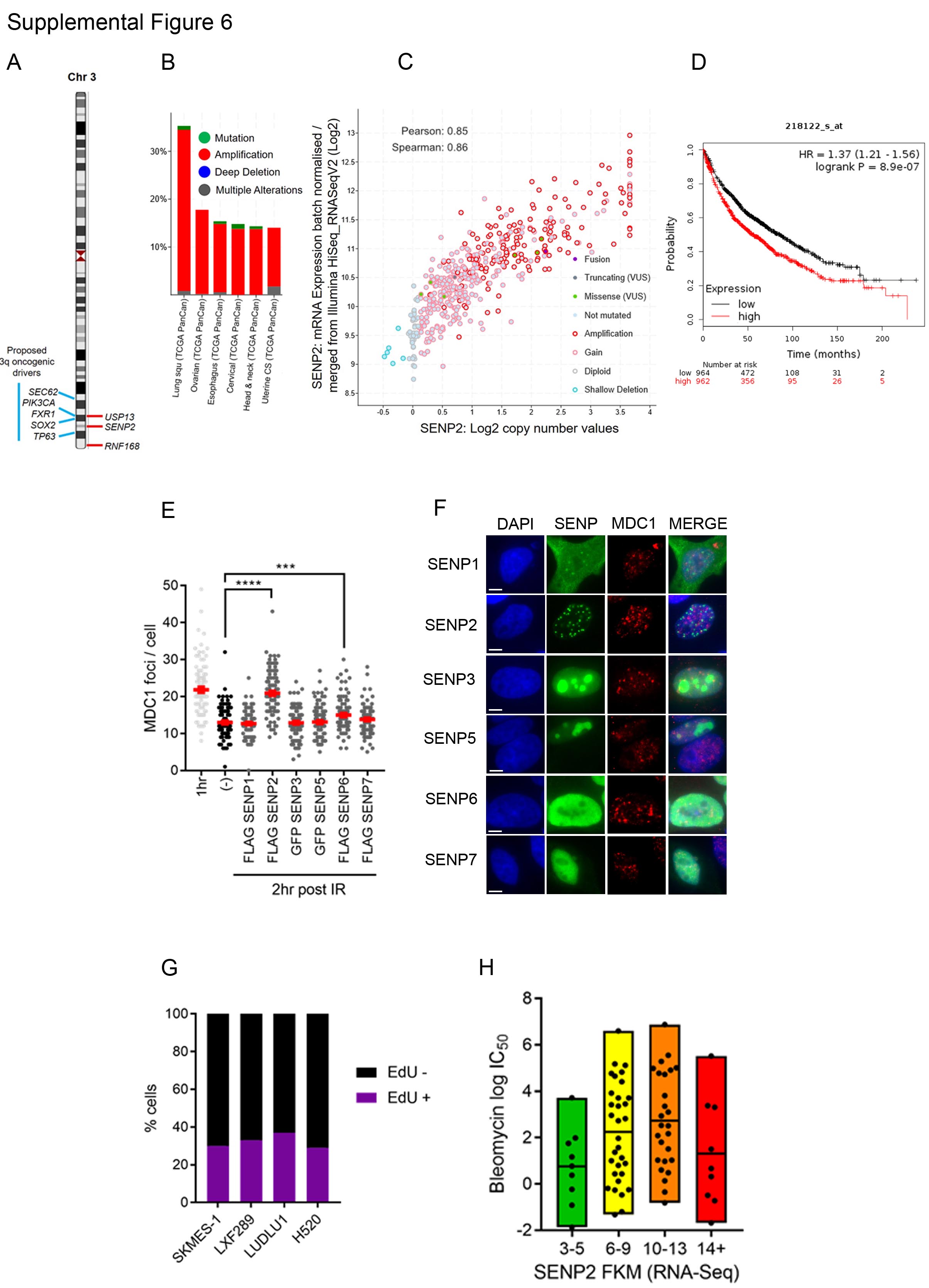

**Supplemental Figure 6. Related to Figure 6.**

**High levels of SENP2 disrupts DSB repair.**

**A**. Illustration of Chromosome 3, indicating the location of *USP13, SENP2 and RNF168*. A selection of candidate oncogenic driver genes within the 3q amplicon are also shown.

**B**. Extent of *SENP2* amplification in the top six 3q amplified cancer types, data is adapted from Cbioportal (August 2018).

**C**. Correlation between SENP2 mRNA and copy number in the LUSC (Lung Squamous Cell Carcinoma) dataset from the TCGA, data is adapted from Cbioportal (August 2018). Pearson correlation co-efficient = 0.85, Spearman co-efficient = 0.86.

**D**. Kaplan-Meier survival plot of overall survival generated via KM Plotter (August 2018) (Gyorffy et al. 2013). Patients (n=1928) were split at the median SENP2 expression as determined by microarray (Affymetrix probe ID 218122_s_at).

**E**. Hela^FlpIn^ cells, transfected with expression constructs for SENP1, SENP2, SENP3, SENP5, SENP6 and SENP7 for 48 hr before exposure to 4 Gy IR. Non transfected cells were either allowed to recover for 1 hour, or 2 hours, and SENP expressing cells were allowed to recover for 2 hours. n=100 Graph shows mean number MDC1 foci per cell, error bars are s.e.m.

**F**. representative images from **E**. illustrating SENP expression. Scale bar = 5μm

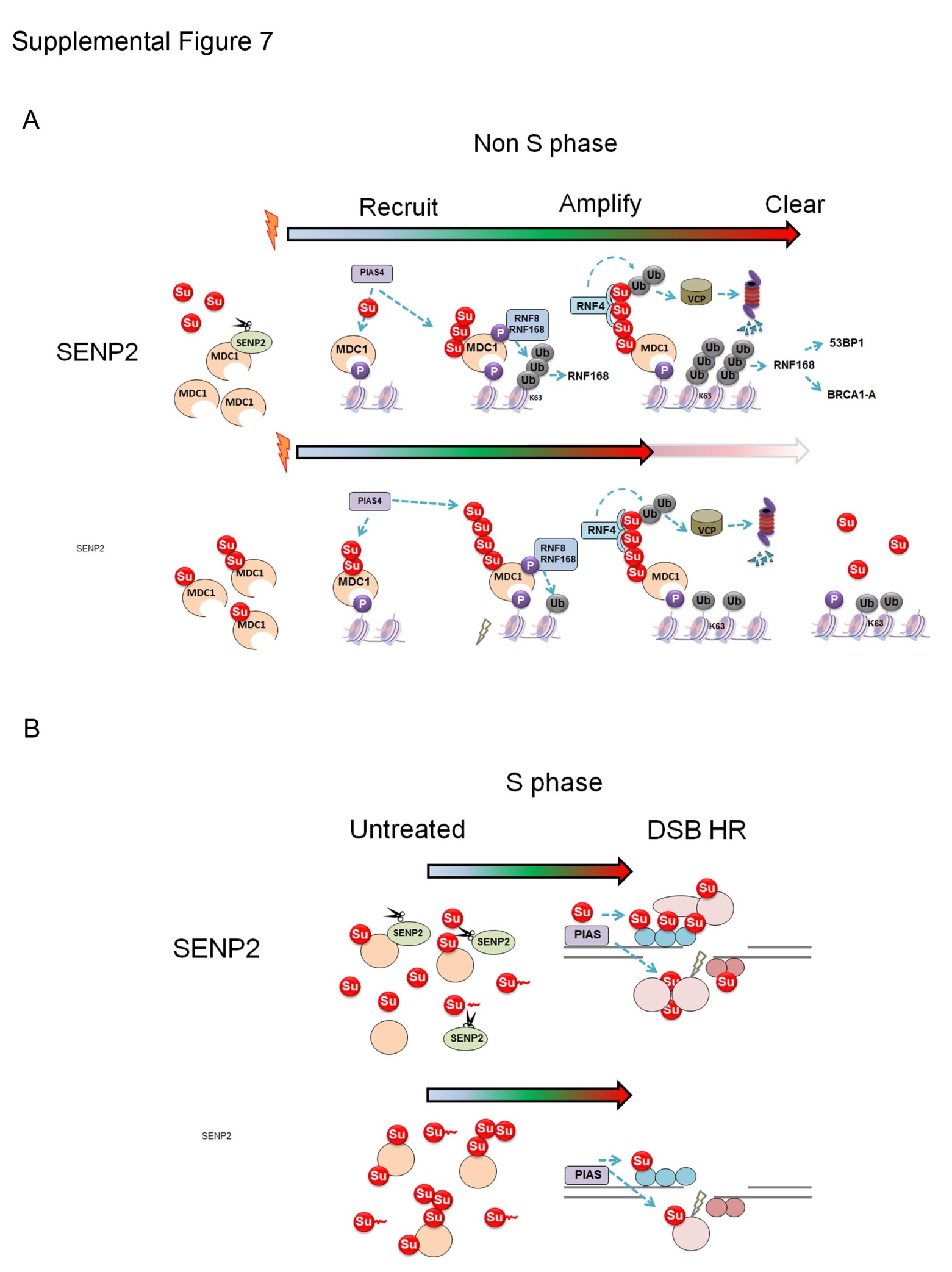

**Supplemental Figure 7.**

**Model of SENP2 action in promoting DSB repair.**

**A**. SENP2 interacts with MDC1 and constitutively cleaves SUMO from it. On damage interaction with SENP2 is lost. MDC1 recruited to chromatin is modified by PIAS4 SUMO E3 ligase and recruits RNF8/RNF168 which lay down K63-Ub marks leading to 53BP1 and BRCA1-A complex recruitment. RNF4 engages with SUMO-MDC1 once sufficient SUMO is conjugated to MDC1, resulting in ubiquitination and VCP engagement.

Without SENP2, MDC1 is recruited to chromatin bearing SUMO moieties and is engaged by RNF4-VCP before sufficient K63-Ub is generated by RNF8/RNF168.

Note although polySUMO of MDC1 is illustrated here for clarity we do not discount that the single SUMO site required encourages multi-mono-SUMOylation that engages RNF4.

**B.** SENP2 cleaves SUMO from cellular conjugates and processes immature SUMO so that on induction of DSBs sufficient SUMO is available for incorporation into multiple interactions required for HR, group modification and or specific SUMO-mediated interactions. Without SENP2 SUMO remains in conjugates and immature SUMO is less efficiently processed so that insufficient SUMO isoform is available for HR.

**Supplemental table 1. Antibodies**

| **Target** | **Host / clonality** | **Vendor** | **Use** | **Catalogue number** | **RRID** |
| --- | --- | --- | --- | --- | --- |
| 53BP1 | Goat polyclonal | R & D Systems | 1:2000 (IF) | AF1877 | AB_2206635 |
| 53BP1 | Rabbit polyclonal | Abcam | 1:2000 (IF) | ab36823 | [AB_722497](http://antibodyregistry.org/AB_722497) |
| ATXN3 | Rabbit polyclonal | Abcam | 1:1000 (WB) | ab96316 | AB_10680570 |
| β-actin | Rabbit polyclonal | Abcam | 1:2000 (WB) | ab8227 | AB_2305186 |
| BRCA1 (MS110) | Mouse monoclonal | Calbiochem | 1:500 (WB) | OP92 | AB_564282 |
| BRCA1 (D9) | Mouse monoclonal | Santa Cruz | 1:500 (IF) | sc6954 | AB_626761 |
| FLAG M2 | Mouse monoclonal | Sigma | 1:2000 (WB/IF) | F1804 | AB_262044 |
| GAPDH | Mouse monoclonal | Abcam | 1:2000 (WB) | ab8245 | AB_2107448 |
| His | Mouse monoclonal | Sigma | 1:2000 (WB) | H1029 | AB_260015 |
| H2AX-pSer139 | Mouse monoclonal | Abcam | 1:2000 (IF) | ab2893 | AB_303388 |
| H2AX-pSer139 | Rabbit polyclonal | Abcam | 1:2000 (IF) | ab22551 | AB_447150 |
| Lamin B1 | Rabbit polyclonal | Abcam | 1:1000 (WB) | ab16048 | AB_443298 |
| MDC1 | Rabbit polyclonal | Abcam | 1:1000 (WB) | ab11169 | AB_297807 |
| MDC1 | Rabbit polyclonal | Bethyl | 1:1000 (WB) | PLA-0016 | AB_203282 |
| MDC1 | Mouse monoclonal | Abcam | 1:1000 (WB) | ab50003 | AB_881103 |
| MYC | Mouse monoclonal | Abcam | 1:2000 (WB) | ab32 | AB_303599 |
| NUP107 | Rabbit monoclonal | Abcam | 1:1000 (WB) | ab178399 | AB_2620147 |
| NUP153 | Mouse monoclonal | Abcam | 1:1000 (WB) | ab24700 | AB_2154467 |
| PIAS1 | Rabbit monoclonal | Abcam | 1:1000 (WB) | ab109388 | AB_10867435 |
| PIAS4 | Mouse monoclonal | Abcam | 1:500 (WB) | ab211625 |  |
| RAD51 | Rabbit polyclonal | Santa Cruz | 1:200 (IF) | sc8349 | AB_2253533 |
| RFP | Rabbit polyclonal | Abcam | 1:1000 (WB) | ab62341 | AB_945213 |
| RNF168 | Rabbit polyclonal | Calbiochem | 1:1000 (WB) | ABE367 | AB_11205761 |
| RNF4 | Goat polyclonal | R & D Systems | 1:1000 (WB) | AF7964-100 |  |
| RPA70 | Mouse monoclonal | Calbiochem | 1:100 (IF) | NA18 | AB_213121 |
| SENP1 | Rabbit monoclonal | Abcam | 1:1000 (WB) | ab108981 | AB_10862449 |
| SENP2 | Rabbit monoclonal | Abcam | 1:1000 (WB) | ab124724 | AB_10972485 |
| SUMO1 | Rabbit monoclonal | Abcam | 1:1000 (WB) | ab32058 | AB_778173 |
| SUMO1 | Rabbit polyclonal | Santa Cruz | 1:200 (IF) | FL-101 | AB_661458 |
| SUMO2/3 | Mouse monoclonal | Abcam | 1:2000 (WB) | ab32058 | AB_1658424 |
| SUMO2/3 | Rabbit polyclonal | Santa Cruz | 1:200 (IF) | FL-103 | AB_2286894 |
| Ub K63 Apu3 | Rabbit monoclonal | Calbiochem | 1:200 (IF) | 05-1308 | AB_1587580 |
| USP13 | Rabbit polyclonal | Sigma | 1:1000 (WB) | HAP004827 | AB_1080497 |
| Vinculin | Rabbit monoclonal | Abcam | 1:2000 (WB) | ab129002 | AB_11144129 |
| Goat α Mouse AF 488 | Goat polyclonal | Life Tech | 1:2500 (IF) | A11001 | AB_2534069 |
| Goat α Rabbit AF 488 | Goat polyclonal | Life Tech | 1:2500 (IF) | A11008 | AB_143165 |
| Goat α Mouse AF 555 | Goat polyclonal | Life Tech | 1:2500 (IF) | A21422 | AB_141822 |
| Goat α Rabbit AF 555 | Goat polyclonal | Life Tech | 1:2500 (IF) | A21428 | AB_141784 |
| Donkey αMouse AF488 | Donkey polyclonal | Life Tech | 1:2500 (IF) | A21202 | AB_141607 |
| Donkey α Rabbit AF555 | Donkey polyclonal | Life Tech | 1:2500 (IF) | A31572 | AB_162543 |
| Donkey α Goat AF488 | Donkey polyclonal | Life Tech | 1:2500 (IF) | A11055 | AB_2534102 |
| Donkey α Rabbit AF488 | Donkey polyclonal | Life Tech | 1:2500 (IF) | A2106 | AB_141708 |
| Rabbit α Mouse HRP | Rabbit polyclonal | DAKO | 1:5000 (WB) | P0161 | AB_2687969 |
| Swine α Rabbit HRP | Pig polyclonal | DAKO | 1:5000 (WB) | P0217 | AB_2728719 |
| Rabbit α Goat HRP | Rabbit polyclonal | DAKO | 1:5000 (WB) | P0449 | AB_2617143 |
| Goat α Rabbit LC HRP | Goat polyclonal | Millipore | 1:5000 (WB) | AP2009 |  |

**Supplemental table 2. Cell lines**

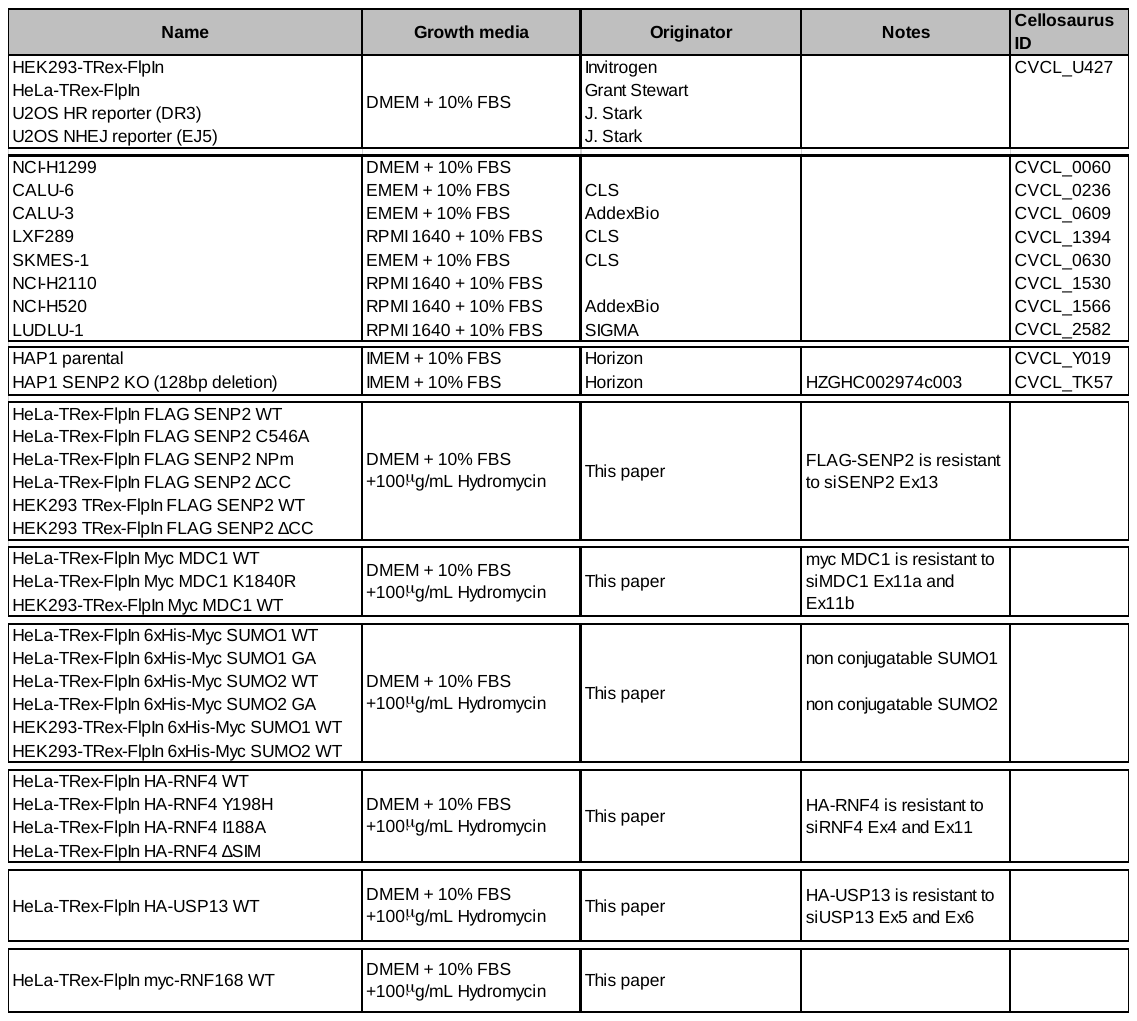

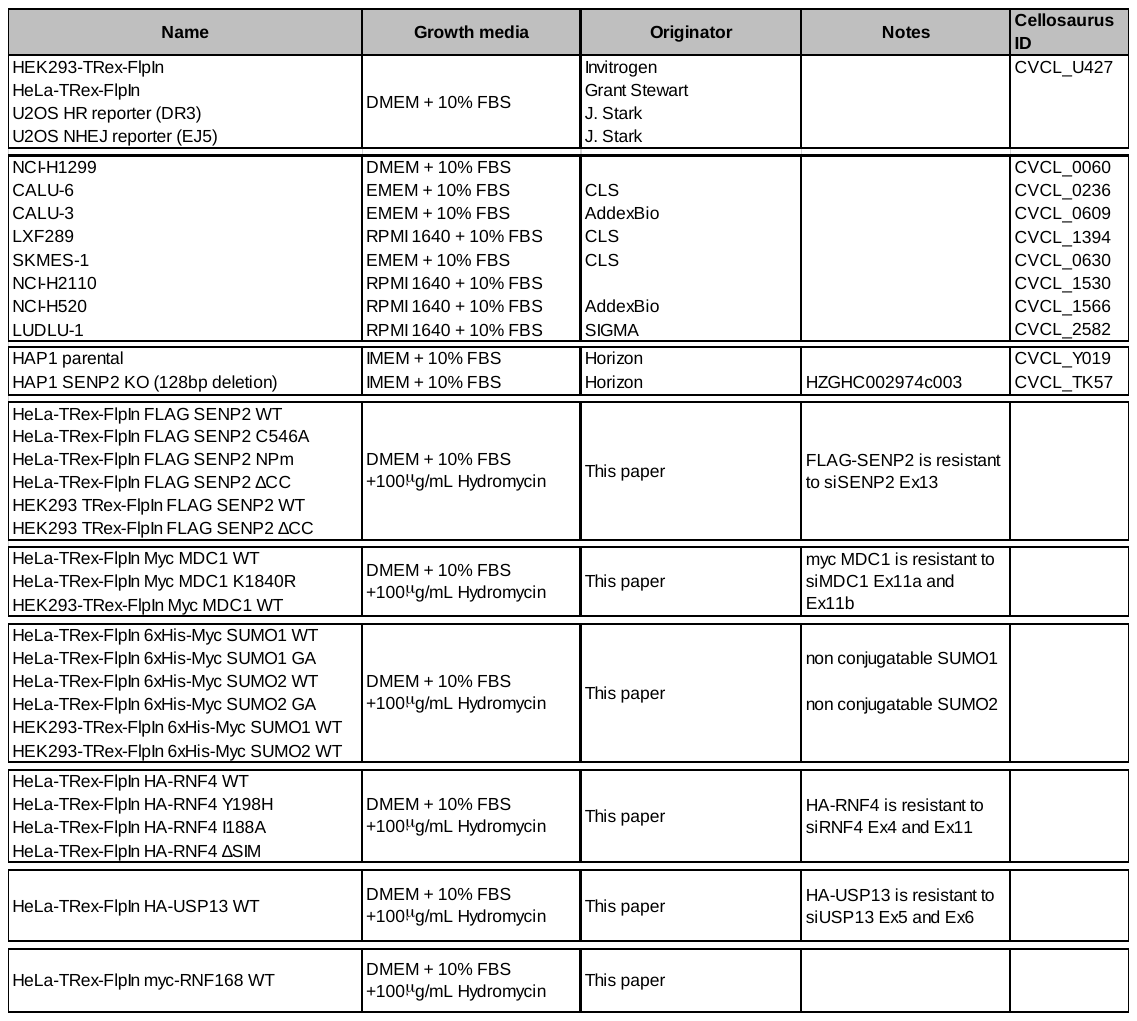

**Supplemental table 3. siRNA**

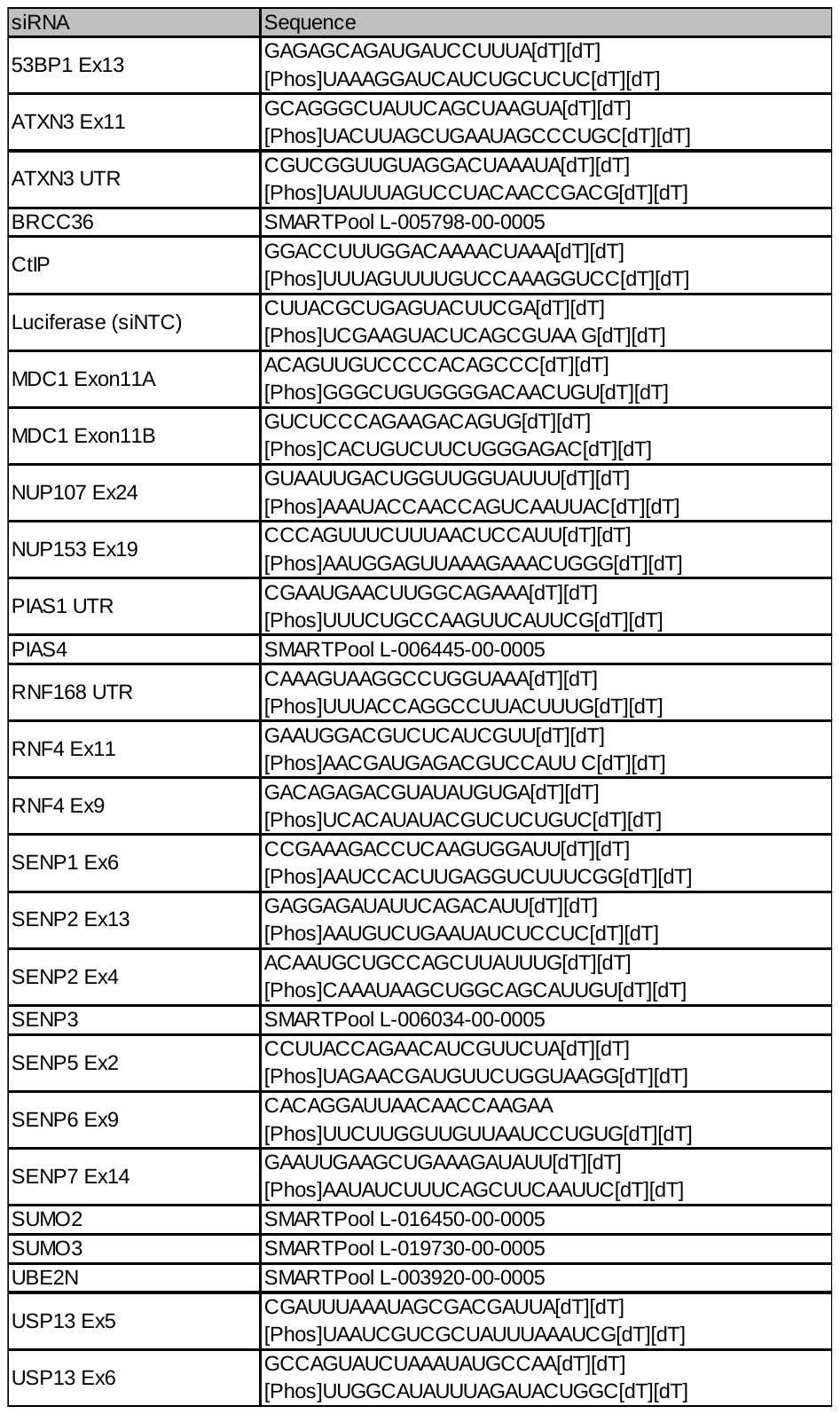

**Supplemental table 4. DNA Oligonucleotides**

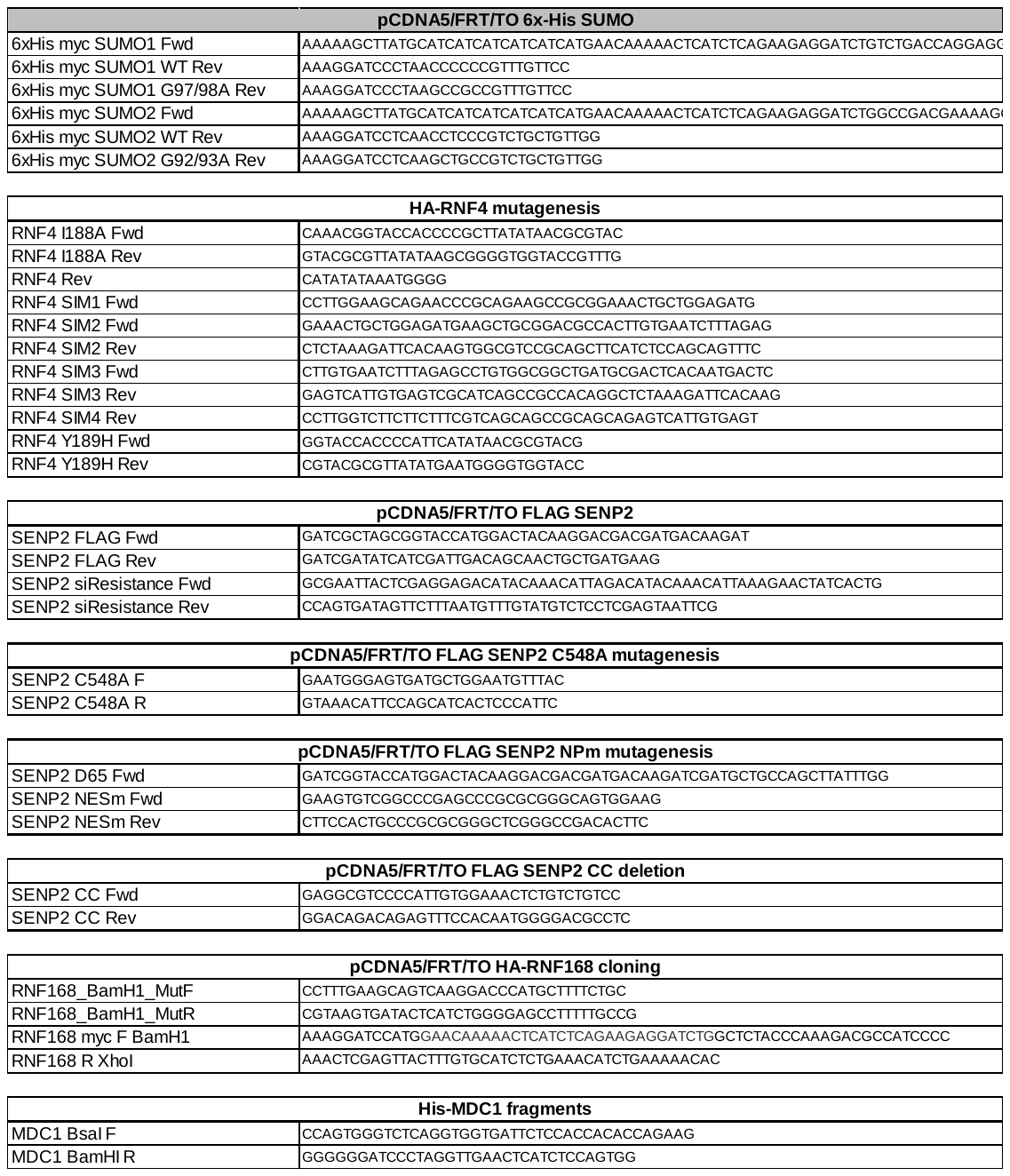

**Supplemental references**

Ashkenazy, H., S. Abadi, E. Martz, O. Chay, I. Mayrose, T. Pupko, and N. Ben-Tal. 2016. 'ConSurf 2016: an improved methodology to estimate and visualize evolutionary conservation in macromolecules', *Nucleic Acids Research*, 44: W344-W50.

Combet, C., C. Blanchet, C. Geourjon, and G. Deleage. 2000. 'NPS@: Network Protein Sequence Analysis', *Trends in Biochemical Sciences*, 25: 147-50.

Gyorffy, B., P. Surowiak, J. Budczies, and A. Lanczky. 2013. 'Online Survival Analysis Software to Assess the Prognostic Value of Biomarkers Using Transcriptomic Data in Non-Small-Cell Lung Cancer', *PLoS ONE*, 8.

Hendriks, I. A., D. Lyon, C. Young, L. J. Jensen, A. C. Vertegaal, and M. L. Nielsen. 2017. 'Site-specific mapping of the human SUMO proteome reveals co-modification with phosphorylation', *Nat Struct Mol Biol*.

Odeh, H. M., E. Coyaud, B. Raught, and M. J. Matunis. 2018. 'The SUMO-Specific Isopeptidase SENP2 is Targeted to Intracellular Membranes via a Predicted N-Terminal Amphipathic alpha-Helix', *Mol Biol Cell*: mbcE17070445.

Rost, B., G. Yachdav, and J. F. Liu. 2004. 'The PredictProtein server', *Nucleic Acids Research*, 32: W321-W26.

Zhang, H., H. Saitoh, and M. J. Matunis. 2002. 'Enzymes of the SUMO modification pathway localize to filaments of the nuclear pore complex', *Mol Cell Biol*, 22: 6498-508.
